# Beyond Kaftrio : mechanistic insights to maximize N1303K-CFTR rescue

**DOI:** 10.1101/2024.02.29.582514

**Authors:** Iwona Pranke, Valeria Capurro, Benoit Chevalier, Emanuela Pesce, Valeria Tomati, Cristina Pastorino, Aurelie Hatton, Saik Urien, Mariateresa Lena, Elise Dréano, Renata Bocciardi, Federico Zara, Stefano Pantano, Vito Terlizzi, Cristina Lucanto, Stefano Costa, Laura Claut, Valeria Daccò, Piercarlo Poli, Massimo Maschio, Benedetta Fabrizzi, Nicole Caporelli, Marco Cipolli, Sonia Volpi, Vincent Jung, Kevin Roger, Frederique Chedevergne, Laure Cosson, Julie Macey, Jean LeBihan, Laurence Weiss, Dominique Grenet, Laurence LeClainche Viala, Benoit Douvry, Bruno Ravoninjatovo, Camille Audousset, Aurélie Tatopoulos, Bénédicte Richaud Thiriez, Melissa Baravalle, Guillaume Thouvenin, Guillaume Labbé, Marie Mittaine, Philippe Reix, Isabelle Durieu, Julie Mankikian, Stéphanie Bui, Mairead Kelly-Aubert, Thao Nguyen–Khoa, Karim Khoukh, Clémence Martin, Chiarra Guerrera, Jennifer Da Silva, Paola di Carli, Carlo Castellani, Federico Cresta, Luis Galietta, Anne Guillemaut, Naim Bouazza, Emmanuelle Girodon, Natacha Remus, Pierre Régis Burgel, Isabelle Sermet-Gaudelus, Alexandre Hinzpeter, Nicoletta Pedemonte

## Abstract

**Introduction:** N1303K is the fourth most frequent Cystic Fibrosis (CF) causing mutation. People with CF (pwCF) clinical status can be improved by Elexacaftor(ELX)/Tezacaftor(TEZ)/Ivacaftor (ETI) combotherapy. We investigated the mechanism underlying N1303K-CFTR rescue.

**Methods:** N1303K-CFTR expression and maturation was evaluated by Western Blot in cell lines and Human Nasal Epithelial Primary Cells (HNECs). Cell surface expression was studied by nanoluciferase complementation assay and TurboID proximity labeling. Functional rescue was tested *in vitro* by YFP-Based Assay and Short Circuit Current.

**Results:** Correction by ELX/TEZ increases N1303K-CFTR amounts, but not its maturation in CFTR-expressing HEK and 16HBEge cell lines and in HNECs. In control conditions, N1303K-CFTR is more distributed at the cell surface and significantly more surface partners are identified in the N1303K-CFTR interactome as compared to F508del-CFTR in HEK cells. ELX/TEZ induces a global stabilization of N1303K-CFTR without favoring its plasma membrane relocation in contrast to F508del-CFTR which is redistributed to the membrane. ETI increases N1303K-CFTR activity in HNECs and can be increased by API co-potentiation with a predicted increase in Forced Expiratory Volume in 1 second (ppFEV_1_) by respectively 13(2)% and 18%(3). This is consistent with a gain in ppFEV1 reported in pwCF carrying the N1303K mutation and additional improvement by API in a patient.

**Conclusion:** These results support the expansion of ETI approval to N1303K mutation but highlight different mechanisms of action than for F508del.

## Introduction

Cystic fibrosis (CF) is a life limiting autosomal genetic disease caused by bi-allelic mutations within the Cystic Fibrosis Transmembrane regulator (*CFTR*) gene. CFTR protein absence or dysfunction and subsequent alterations induce multi-systemic damage resulting in pancreatic insufficiency, chronic bronchopathy, and abnormally salty sweat (1,2).

The quality of life of people with CF (pwCF) was greatly improved by CFTR modulators, e.g; small molecules targeting CFTR (1,3). The most frequent variant, p.Phe508del, (F508del hereafter) alters protein maturation and function (2,3). It can be rescued by combining correctors (VX-445, Elexacaftor, ELX, and VX-661, Tezacaftor, TEZ) that favor maturation and a potentiator (VX-770, Ivacaftor, IVA) that increases channel activity. ELX/TEZ/IVA (ETI hereafter) greatly improves the clinical status of patients carrying F508del on at least one allele (4). Moreover, *in vitro* data in Fischer Rat Thyroid (FRT) cells expressing other *CFTR* variants supported the expansion of ETI approval to 177 other mutations in the USA (5). p.Asn1303Lys (N1303K hereafter) represents the fourth most frequent mutation in pwCF with a relatively high allelic frequency in Italy (5.46%) and France (7.9%) (6,7). N1303K is located within the second nucleotide binding domain (NBD2), causes protein misfolding and displays a modest correction by ETI in different cell lines (8,9,10). However, experiments in FRT cells reported that ETI restored CFTR function to 9.4% of the Wild Type (WT), approaching the 10% threshold considered predictive of clinical efficiency (11,12). Importantly, N1303K-CFTR rescue was also observed in patient-derived primary cell models (13–16). Results of a placebo-controlled clinical study, as well as registry and real-world data provided evidence that pwCF carrying N1303K in the absence of F508del responded to ETI and support the request to EMA of ETI label expansion for pwCF carrying this mutation (14,17,18,19,20,21).

From a mechanistic point of view, the defects associated with N1303K-CFTR appear distinct from F508del-CFTR. N1303K-CFTR degradation has been proposed to involve mainly the autophagy pathway whereas F508del-CFTR is mainly degraded by the proteasome (22). N1303K perturbs the correct alignment of ATP on NBD2, suggesting an uncoupling between NBDs and Transmembrane Domains (TMDs) while F508del induces a correct dimerization of the NBDs but results in a loss of contact between NBD1 and Intra Cellular Loops (ICLs) in the second TMD (10,23). Importantly, it has been reported that N1303K-CFTR activity is improved by IVA, even in the absence of correctors with a further enhancement by co-potentiation with Apigenin (API) (10, 24, 25). This observation, shown in FRT cells, was recently confirmed in rectal organoids where N1303K-CFTR rescue was maximized by the quadruple combination ELX/TEZ/IVA and API (16). We investigated mechanisms underlying N1303K-CFTR rescue in respiratory cells and provide experimental and clinical data supporting the expansion of the label of ETI for N1303K and potential repurposing of API as co-therapy.

## MATERIAL AND METHODS

### Cell models

CFF-16HBEge cell lines expressing N1303K, F508del, G542X were provided by the Cystic Fibrosis Foundation Therapeutics (26). NCL-SG3 human eccrine sweat gland cell line was provided by Thibault Kervarrec (Faculté de Pharmacie, Tours, France) (27). Human Nasal Epithelial cells (HNECs) were obtained by nasal brushing and cultured as previously reported (28–30).

### CFTR expression assays

CFTR expression level and profile were assessed by Western Blot (28–30). CFTR cell surface expression was quantified using the nanoluciferase complementation assay in HEK293 cells (31).

### Ussing chamber studies

Culture inserts with differentiated HNECs were mounted in Ussing chambers. Short-circuit current (Isc) was measured after short-circuiting the trans-epithelial ion flux with a voltage-clamp (14, 28–30). The following activators or inhibitors were added sequentially: amiloride (10μM) at the apical side to block sodium reabsorption; CPT-cAMP (100μM) or alternatively Forskolin/IBMX 10µM/100µM at both apical and basolateral sides to activate CFTR; IVA (1μM) at the apical side to potentiate the channel gate; API (25μM) at the apical side to co-potentiate IVA; and the CFTR specific inhibitor-172 (Inh-172) (20μM) at the apical side (all from Sigma Aldrich).

Isc change after stimulation with cAMP agonists and its inhibition by Inh-172(ΔIsc_inh-172_) served as an index of CFTR function. Only samples with a transepithelial resistance above 300 Ω*cm^2^ were analyzed.

### YFP-Based Assay for CFTR Activity

CFBE41o-cells stably expressing the halide-sensitive yellow fluorescent protein (HS-YFP) were transfected using Lipofectamine 2000 with vectors encoding WT, N1303K or F508del-CFTR (28–30). YFP quenching rate was used to assess CFTR activity.

### CFTR interactome

The interactome of WT, F508del and N1303K-CFTR was assessed in HEK293 cells using the TurboID proximity labeling approach, as previously described (32).

### Patients

All patients carried the N1303K variant on at least 1 allele, in *trans* with a minimal function (MF) variant non rescuable by ETI. Thirty-three Italian patients (Ethics Committee of the Istituto Giannina Gaslini; CER 28/2020, 04/04/2020) provided HNECs by nasal brushing. Twenty-eight French patients entered a compassionate program and were evaluated at baseline and at 1 to 2 months of ETI for Forced Expiratory Volume, expressed in percentage predicted (ppFEV_1_) and sweat chloride concentration (mmol/L) completed by β-adrenergic secretion in selected patients (Institutional Review Board of the *Société de Pneumologie de Langue Française* (#2020–003)(14, 18, 33). Extensive genetic screening was performed in non-responder patients (e.g ; less than 5% improvement in ppFEV_1_) or patients who experienced a respiratory degradation after initial improvement, including next generation sequencing to search for *CFTR* complex alleles and variants of genes that were shown to modify airway surface liquid homeostasis (*TMEM16A, SLC26A4, ATP12A, and SCNN1A/B/G)*, or respiratory status such as *SERPINA1* gene (34).

Additional information for all these assays are described in **Supplemental Material**.

### Statistics

Statistical analyses were performed with Statview or R. Comparisons were made by two-tailed Student’s t-test, one-way analysis of variance (ANOVA) with Tukey’s or Dunett’s post-test, or Wilcoxon and Mann-Whitney nonparametric tests and Repeated Measures ANOVA for paired samples. Modeling of CFTR half-life according to mutant is described in **Supplemental Material**. p<0.05 was considered significant.

## RESULTS

### 1. ELX and TEZ combination increases N1303K-CFTR expression but not its maturation

In transiently transfected HEK293 cells, N1303K-CFTR was detected in control conditions as a mature fully glycosylated (band C, at ∼180 kD) and an immature core-glycosylated CFTR (band B at ∼150 kD), (**Figure 1a**). ELX/TEZ correctors increased both band B and band C of N1303K-CFTR without modifying its maturation, as assessed by the unchanged C/(C+B) ratio. This contrasted with F508del-CFTR where only band B could be detected at baseline while ELX/TEZ doubled the maturation ratio, consistent with the conversion of immature into mature F508del-CFTR (**Figure 1a**). A similar pattern was present in 16HBEge cell lines (**Figure 1b**). In HNECs, N1303K-CFTR was mainly detected as a band B with a faint band C. ELX/TEZ treatment increased both band B and band C, while an obvious B to C switch was detected in F508del HNECs (**Figure 1c and 1d).** This pattern of response was not tissue dependent, as shown in the sweat gland NCL-SG3 and the lung epithelial 16HBEge-G542X-CFTR cell line, not expressing CFTR, both transiently transfected with plasmids encoding N1303K-CFTR or F508del-CFTR (**Figure Supplemental 1a and 1b**). In both cell lines, N1303K-CFTR band C was also present in control condition and correctors increased both band B and C but not the C/(C+B) ratio. In contrast, in F508del-CFTR cells, ELX/TEZ induced a clear switch to the mature form. This profile was not changed by autophagy inhibition both in those cell lines (**Figure Supplemental 2a**) and in HNECs (**Figure Supplemental 2b**).

**Figure 1.**
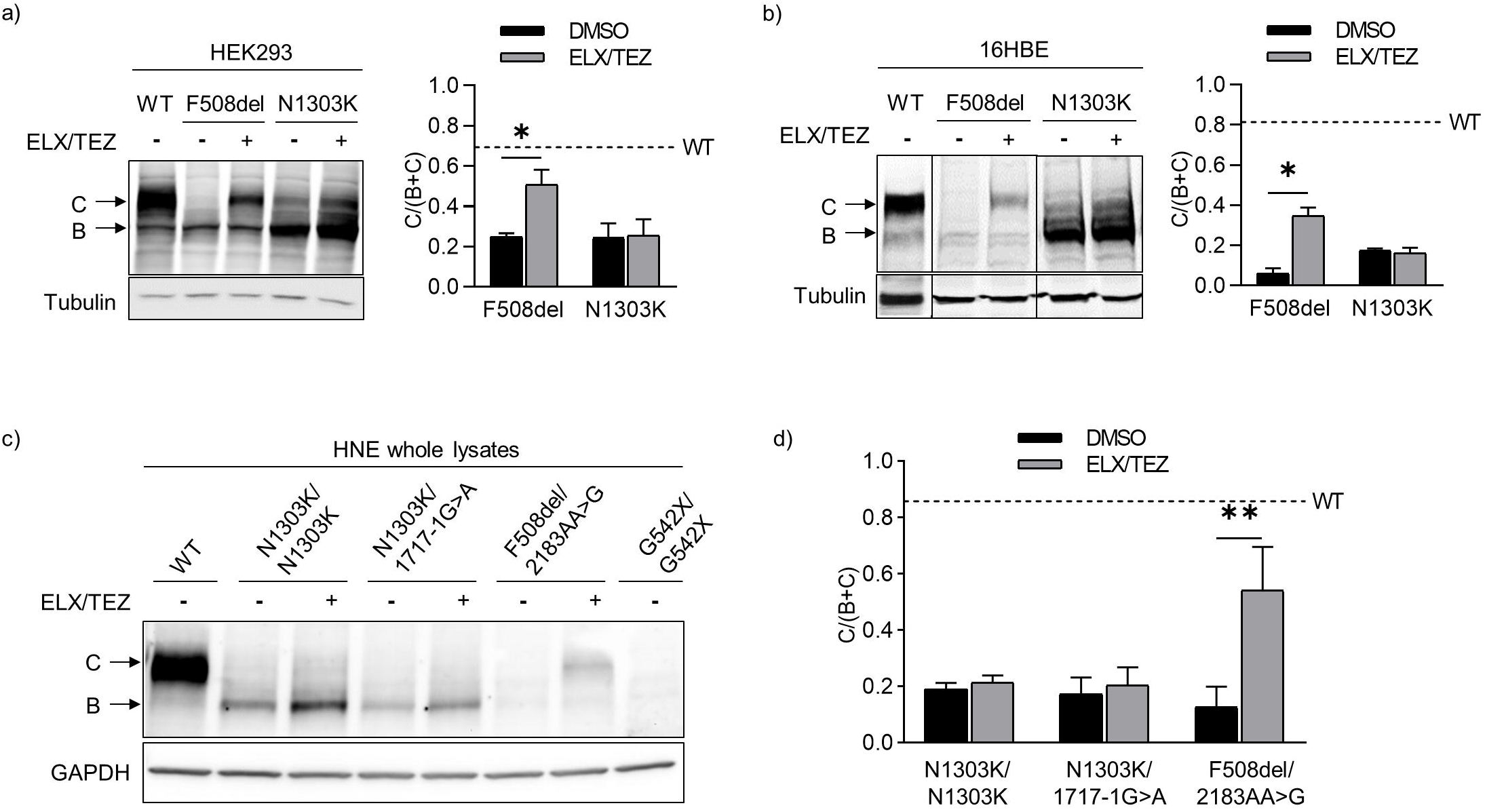
Maturation and correction of N1303K-CFTR. Representative Western blot images and corresponding quantifications for CFTR-N1303K, CFTR-WT and CFTR-F508del incubated for 24 hours with DMSO (vehicle) or correctors elexacaftor/tezacaftor (ELX/TEZ at 3 µM each) from a) transiently transfected HEK293 cells and b) 16HBEge-N1303K-CFTR cells and c) Human Nasal Epithelial primary cells (HNECs). HNECs were obtained from: non-CF donor (WT; donor ID: Ctr069), homozygote N1303K (donor ID: ME084), heterozygote N1303K/1717-1G>A (donor ID: GE011), heterozygote F508del/2183AA>G (donor ID: AN238) and homozygote G542X (donor ID: GE220). For a), b) and c), bar graphs on the right panel correspond to the quantification of CFTR maturation expressed as a C/(C + B) band ratio for each different cell type. As a reference, the dotted bar on the graph corresponds to the ratio for WT. Data are presented as mean ± standard deviation (SD) from a minimum of three independent experiments. * for p<0.05 and ** for p<0.01.

As N1303K-CFTR band B levels were greater as compared to that of F508del-CFTR **(Figure 1a, 1b, 1c)**, we evaluated protein stability in HEK293 cells. Under protein synthesis inhibition by cycloheximide, in control conditions, N1303K-CFTR decreased significantly less over time than F508del-CFTR with a calculated half-life of 11±1.3h *versus* 2±0.4h for F508del-CFTR (p<0.0001). ELX/TEZ significantly increased the half-life of N1303K-CFTR (19±2h) and F508del-CFTR (6±0.8h) (p<0.0001 for both) (**Figure 2a)**. ELX/TEZ slowed down N1303K-CFTR degradation significantly more than that of F508del-CFTR (p<0.0001) (**Figure 2b and Figure Supplemental 3a**).

**Figure 2.**
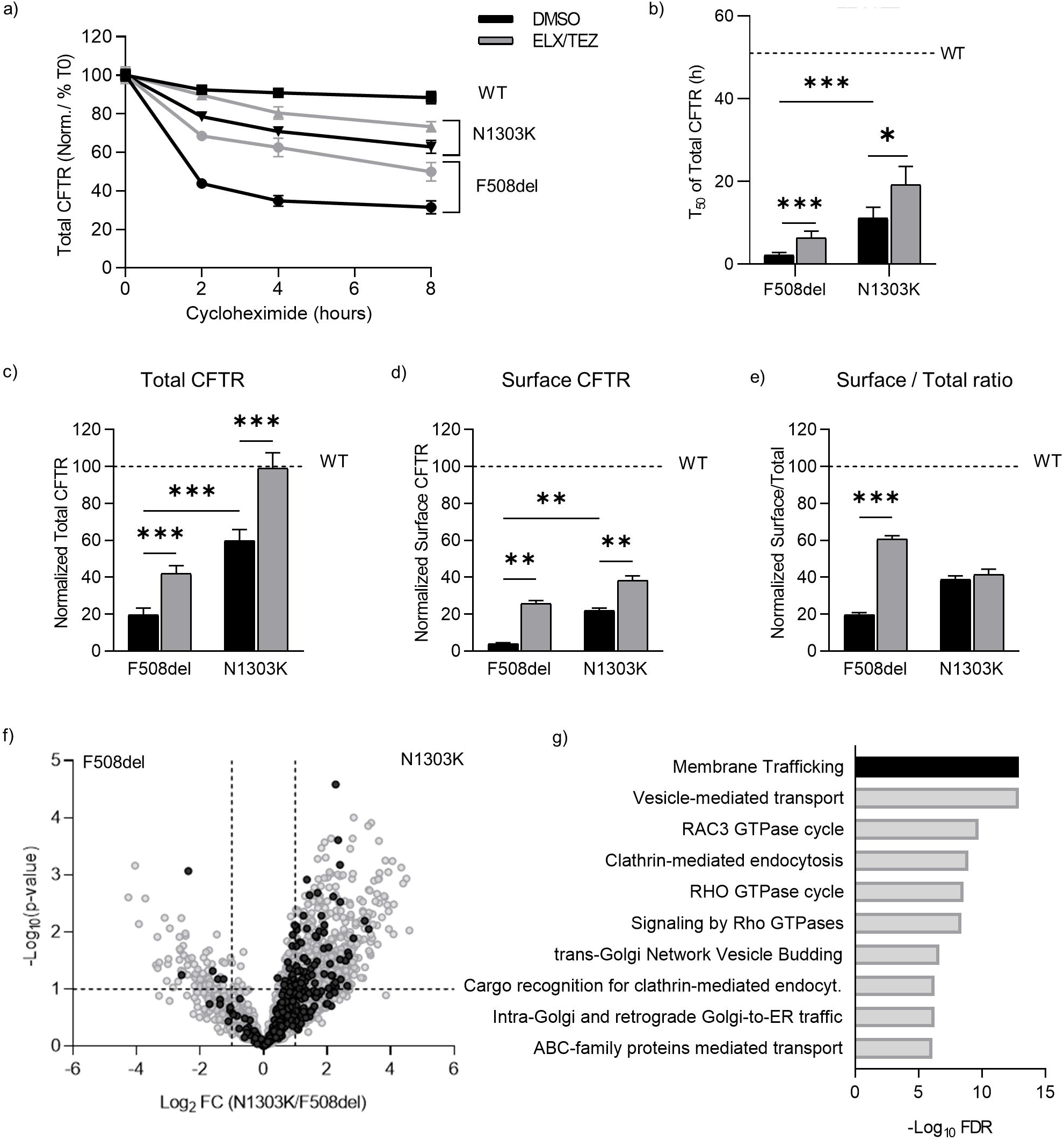
Stability and Surface expression of N1303K-CFTR. a) Total CFTR luminescent signal quantified in HEK293 cells transfected with HiBiT-WT, F508del, or N1303K-CFTR and treated with cycloheximide (CHX) for different timespans (0-8h) and then incubated for 24 hours with DMSO or correctors elexacaftor/tezacaftor (ELX/TEZ at 3 µM each). Luminescent signals were normalized to HiBiT-WT-CFTR in DMSO conditions and data are presented as mean ± SEM (n=4). b) Estimated CFTR half-life obtained from distribution plots derived from modeling total CFTR stability under CHX (T_50_ of Total CFTR) overlayed with the observed data (points). Data are presented as mean ± SEM. ** for p<0.01 and *** for p<0.001 (unpaired t-test, n=4) c-e) Nanoluciferase complementation assay quantification of c) Total CFTR, d) Surface, and e) Surface/Total ratio of CFTR in HEK293 cells transfected with HiBiT-WT, F508del or N1303K-CFTR and treated for 24 hours with either DMSO or correctors (ELX/TEZ, 3 µM each) when indicated. Data are presented as mean ± SEM. ** for p<0.01 and *** for p<0.001 (unpaired t-test, n=5) f,g) N1303K-CFTR interactome obtained by TurboID proximity labelling. f)Volcano plot illustrating the Fold Change between TurboID-N1303K-CFTR and TurboID-F508del-CFTR proteins (Log_2_FC). Proteins referenced in “Membrane trafficking” in g) highlighted in black. g) Reactome Over-Representation Analysis of proteins enriched in TurboID-N1303K-CFTR samples as compared to TurboID-F508del-CFTR. Bar plot showing the top over-represented pathways ordered by -log_10_ estimated False Discovery Rate (FDR).

In control conditions, nanoluciferase complementation assay detected a total CFTR signal significantly greater in N1303K *versus* F508del-CFTR HEK293 cells (**Figure 2c**) and identified N1303K-CFTR at the cell surface while F508del-CFTR was almost absent (**Figure 2d**). ELX/TEZ increased CFTR total and surface expression in the same proportion for N1303K and F508del-CFTR (**Figure 2c** and **2d**). Interestingly, when normalized to the total amount, F508del-CFTR exhibited a significant 3-fold increase of the cell surface ratio while that of N1303K-CFTR was unchanged (**Figure 2e**), providing evidence of a relocalization of F508del-CFTR to the membrane in contrast to N1303K-CFTR. L206W, another class II processing mutation exhibited the same surface profile as F508del-CFTR (**Figure Supplemental 4**). TurboID proximity labeling showed that samples from N1303K-CFTR were enriched in proteins involved in membrane trafficking (**Figure 2f**), vesicle mediated transport and endocytosis as compared to F508del-CFTR (**Figure 2g**). This was confirmed by targeted analysis of known CFTR partners (**Figure Supplemental 5** and **Supplemental Results**)

### 2. N1303K-CFTR activity is increased by the ETI/API combination in epithelial cell lines

In CFBE41o-cells stably expressing HS-YFP, F508del-CFTR correction with ELX/TEZ induced a major increase in CFTR activity (p<0.0005) with a further increase by IVA (p<0.005) but not by IVA/API combination (**Figure 3a**). CFBE-N1303K cells also displayed a significant improvement in CFTR activity under ELX/TEZ/IVA (p<0.05) but in contrast, API increased significantly CFTR activity (p<0.0005) (**Figure 3a**).

**Figure 3.**
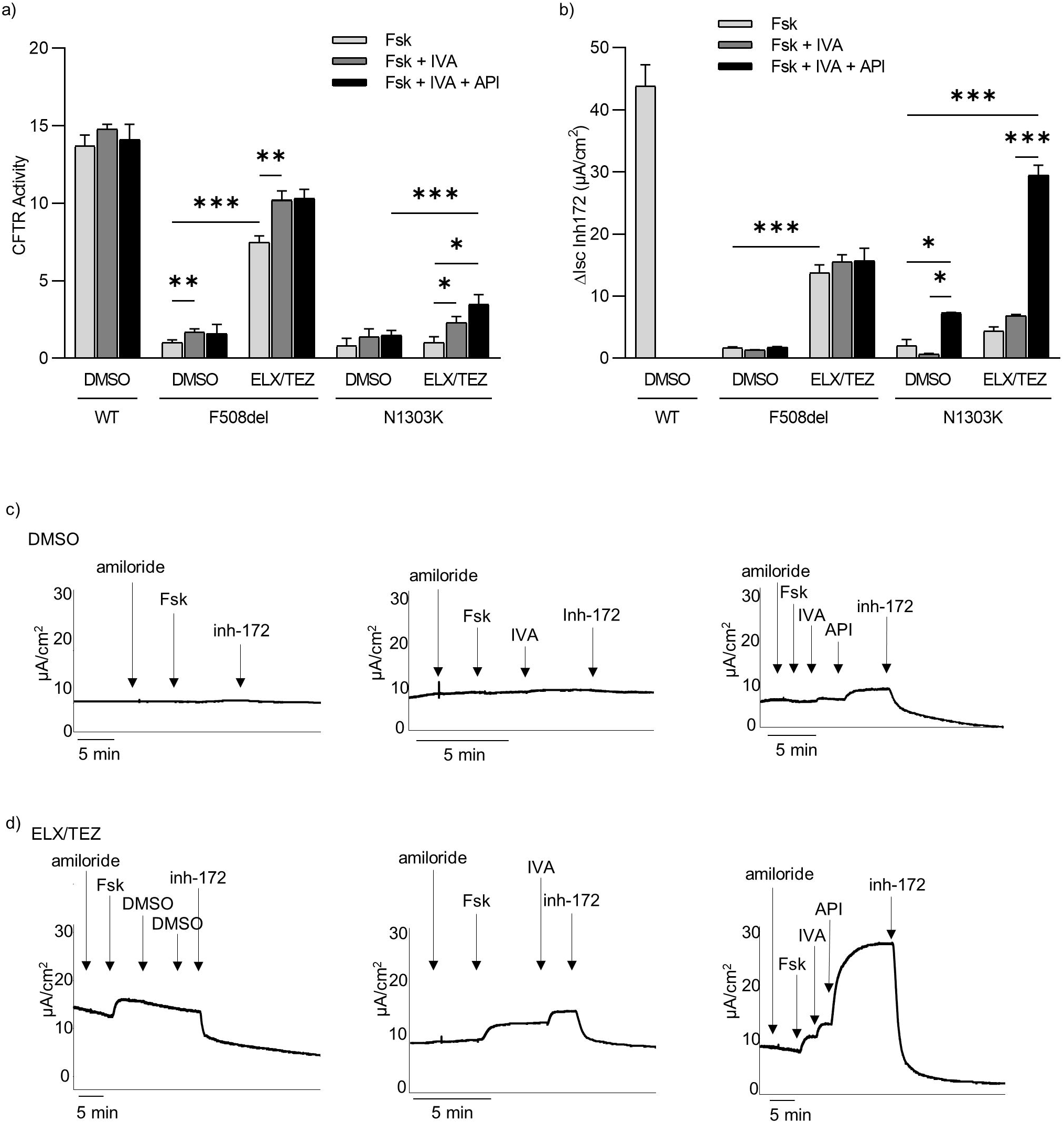
N1303K-CFTR activity potentiation by Ivacaftor and Apigenin in Tezacafor/Elexacaftor corrected CFBE41o- and 16HBEo-cell lines. a) CFTR-N1303K activity in CFBE41o-cells stably expressing HS-YFP. The bar graph shows the activity of N1303K-CFTR transiently expressed in CFBE41o-cells. CFTR activity was determined as a function of the YFP quenching rate following iodide influx elicited by acute incubation of Forskolin (Fsk) (20 µM) (light grey), Fsk + Ivacaftor (IVA) (1 µM) (medium grey), or Fsk + IVA + Apigenin (API) (25 µM) (dark) in cells treated for 24 h with DMSO (vehicle) or with tezacaftor (TEZ 10 µM) combined with elexacaftor (ELX 3 µM). Data from cells transiently expressing WT-CFTR followed by treatment with DMSO and F508del-CFTR followed by treatment with DMSO and ELX/TEZ are also shown for comparison. Data expressed as mean ± standard deviation (SD) from a minimum of three independent experiments. Comparison by unpaired t-test, * for p<0.05, ** p<0.005, *** p<0.0005 b) CFTR activity in 16HBEge-N1303K-CFTR cell lines assessed by Short Circuit Current (Isc). CFTR activation elicited by acute addition in the Ussing chamber of Fsk (10 µM) (light grey), or Fsk+ IVA (10 µM) (medium grey), or Fsk + IVA + API (20 µM) (dark), in cells treated for 48 h with DMSO (vehicle), or with TEZ/ELX (3 µM each). Data from cells transiently expressing either WT-CFTR followed by treatment with DMSO or expressing F508del-CFTR followed by treatment with DMSO or ELX/TEZ are also shown for comparison. Experiments were performed three times and results presented as mean+/-SD. 2-way ANOVA statistical test, * p<0.05, *** p<0.0005. c) and d) Representative short-circuit currents obtained in 16HBEge-N1303K-CFTR cells. Cells were incubated for 48 hours with c) DMSO or d) ELX/TEZ (3 µM each). During the recordings, the epithelia were sequentially treated with amiloride (100 μM), Fsk and IBMX (10µM and 100 µM respectively), IVA (10 µM), API (20 µM), CFTR inhibitor inh-172 (inh-172, 10µM).

In 16HBEge-N1303K-CFTR cells IVA/API combination increased significantly CFTR activity in non-corrected cells (p<0.05) while IVA alone was ineffective (**Figure 3b, 3c**). ELX/TEZ increased activity of N1303K-CFTR and F508del-CFTR to respectively 10.1(2)% (p>0.05) and 31.3(5)% (p<0.0005) of the WT (**Figure 3b, 3d**). IVA combination to ELX/TEZ (ETI) did not modify the chloride secretion in N1303K or F508del-CFTR cells, but its combination with API increased CFTR activity of N1303K-CFTR up to 67.2(6)% of the WT (p<0.0001 *versus* ETI) while this co-potentiation was not observed in 16HBE-F508del-CFTR corrected cells (**Figure 3b, 3d**).

### 3. N1303K-CFTR activity is increased by API co-potentiation in nasal epithelial cells

CFTR activity was studied in HNECs from 28 patients carrying N1303K in *trans* with a MF mutation and of 5 N1303K homozygous patients (**Table 1 Supplemental**). In DMSO treated cells, average N1303K-CFTR function displayed a large interindividual variability as shown in **Figure 4a**. CFTR activity in these non-corrected cells was not significantly increased by IVA, while addition of API after IVA enhanced CFTR activity to 8(5)% of the WT (p<0.0001 *versus* baseline) **(Figure 4a and 4b)**. In ELX/TEZ-corrected HNECs, CFTR activity reached 10.2(6)% of the WT in the presence of IVA alone (ETI combination) (p<0.0001 *versus* baseline) and was further increased up to 14.5(7)% of the normal when API was added to co-potentiate IVA (p<0.0001 *versus* baseline, p<0.0001 *versus* ETI) **(Figure 4a and 4b)**. This level of rescue corresponded in our previously published data set to a predicted absolute gain in ppFEV1 of 12.8(6.2) %, for ETI and 16.6(5.8) % for ETI API combination (**Figure Supplemental 6**) (14). The level of correction remained however variable ranging for ETI between 2.2 to 36.5% and for ETI/API between 4.6% and 40.9% with respective corresponding ppFEV1 gain between 3.7% to 27.3% for ETI and 7.3% and 28% for ETI/API. This was not related to basal activity or the genotype (e.g; 1 or 2 N1303K alleles). This variability was not observed in HNECs of patients carrying F508del, L206W or R334W (**Figure 4c**).

**Figure 4.**
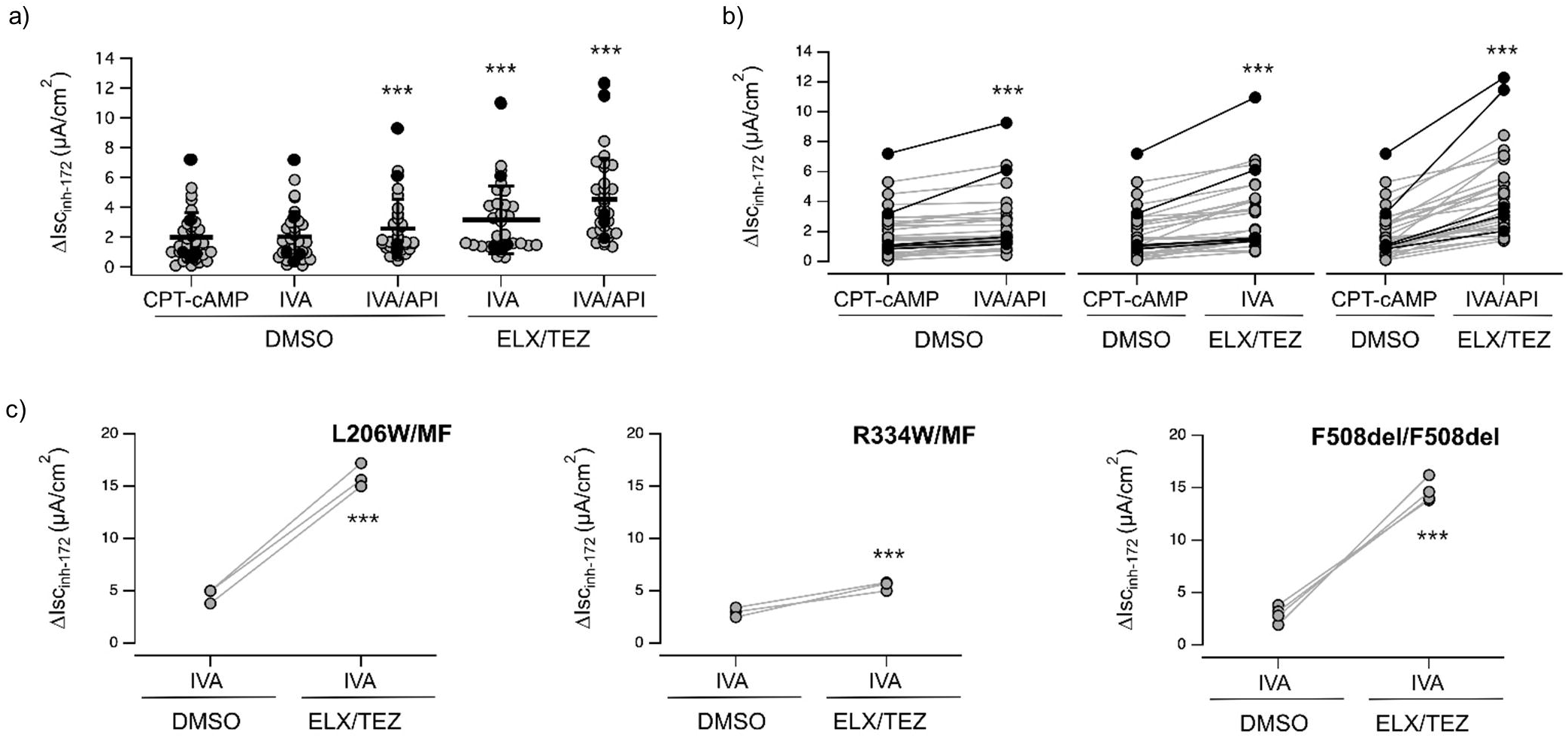
N1303K-CFTR activity potentiation by Ivacaftor and Apigenin in Tezacafor/Elexacaftor corrected Human Epithelial Nasal Epithelial cells. CFTR activity quantified with Short-circuit current technique on nasal epithelial cells treated for 24 hrs with vehicle (DMSO) or ELX/TEZ combination (3 µM/10 µM). During the recordings, the epithelia were sequentially treated with amiloride (10 μM; added on the apical side), CPT-cAMP (100 μM; added on both apical and basolateral sides), IVA (1 μM; apical side), API (25 μM; apical side), and the CFTR inhibitor-172 (inh-172; 20 μM; apical side). Data reported are the average amplitude of the current blocked by 20 μM inh-172 (ΔIscinh-172). a) Scatter dot plots showing experiments performed on twenty-eight subjects compound heterozygous for N1303K and a minimal function (MF), non-rescuable variant (N1303K/MF) and five homozygous for N1303K (N1303K/N1303K) under different conditions. b) Connected dot plots of patients shown in a) c) Connected dot plots obtained during experiments performed with the short-circuit current technique on nasal epithelial cells derived from three patients compound heterozygous for L206W and a MF, non-rescuable CFTR variant (L206W/MF - left panel), three patients compound heterozygous for R334W and a MF, non-rescuable CFTR variant (R334W/MF - middle panel) and four patients homozygous for F508del (F508del/F508del - right panel).

As a whole, N1303K-CFTR-mediated currents displayed four different response patterns **(i)** no or negligible effects of modulators (6 patients) (**Figure 5a)**; **(ii)** no CFTR activity in control conditions but significant response to ETI enhanced by API (11 patients) (**Figure 5b)**; **(iii)** response upon addition of IVA/API in non-corrected cells maximized in ELX/TEZ corrected cells (10 patients) (**Figure 5c); (iv)** CFTR activity rescue only in the quadruple combination of ETI/API (6 patients) (**Figure 5d).**

**Figure 5.**
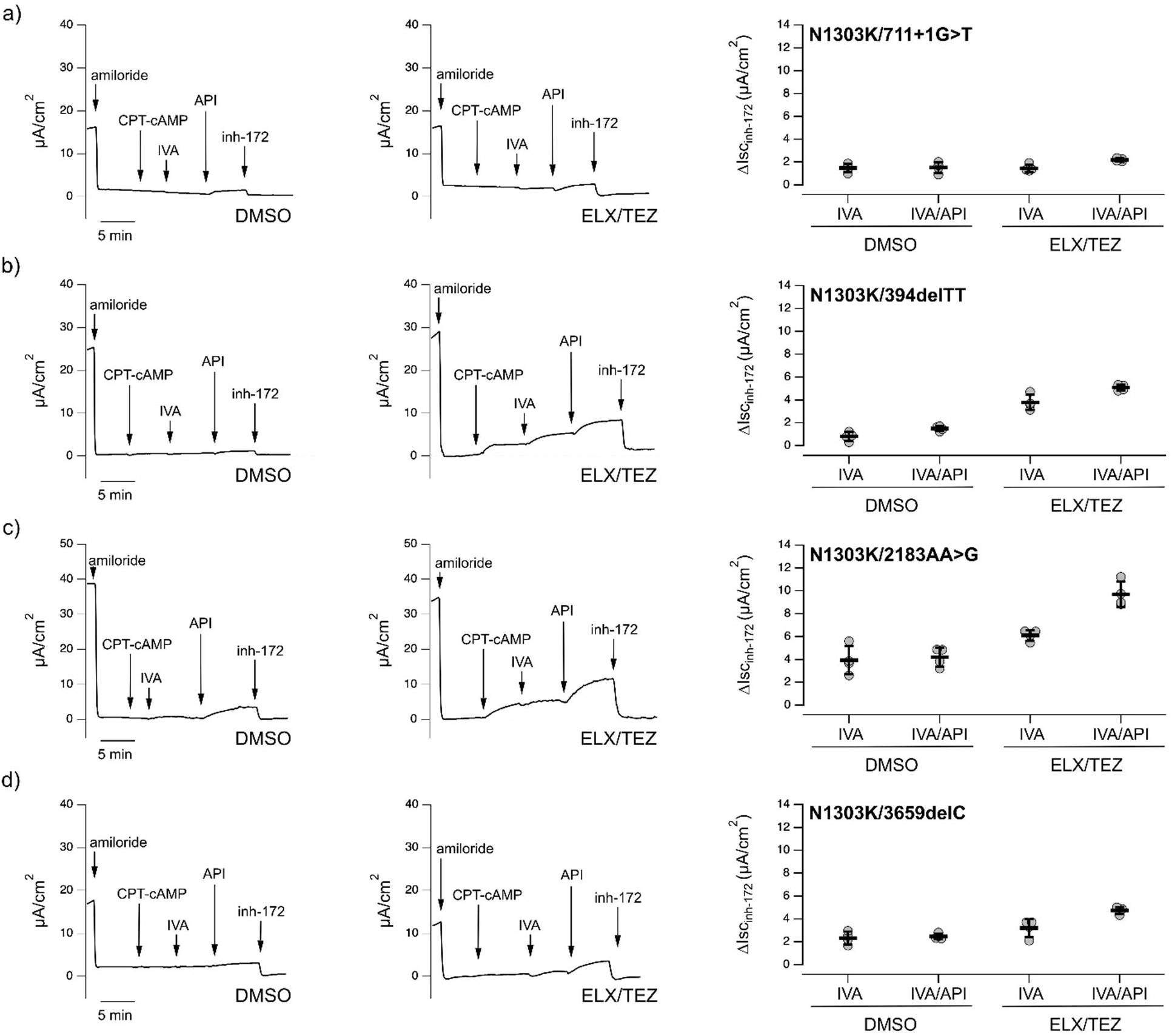
Profiles of N1303K-CFTR activity rescue by modulators in Human Epithelial Nasal Epithelial cells. Representative traces on the left panel of the effect of vehicle (DMSO), or the elexacaftor/tezacaftor (ELX, 3 µM / TEZ, 10 µM) combination on nasal epithelial cells with the short-circuit-current technique. During the recordings, the epithelia were sequentially treated (as indicated by downward arrows) with amiloride (10 μM; added on the apical side), CPT-cAMP (100 μM; added on both apical and basolateral sides), ivacaftor (IVA, 1 μM; apical side), apigenin (API, 25 μM; apical side where indicated), and the CFTR inhibitor-172 (inh-172; 20 μM; apical side). The dashed line indicates zero current level. Right panel: scatter dot plot showing the summary of results. Data reported are the amplitude of the current blocked by 20 μM inh-172 (ΔIscinh-172). For each experimental condition the number of biological replicates were n = 4-6. Short-circuit current performed on: a) N1303K/711+1G>A nasal epithelial cells (derived from donor ID: GE156) b) N1303K/394delTT nasal epithelial cells (derived from donor ID: GE010). c) N1303K/2183AA>G nasal epithelial cells (derived from donor ID: GE132). d) N1303K/3659delC nasal epithelial cells (derived from donor ID: ME119).

### 4. Effect of ETI is variable in N1303K patients and can be improved by API

**Supplemental table 2** shows data reported in 13 N1303K/N1303K and 23 N1303K/MF pwCF in France (14, 18, 21), combined with other published cases in Israel and Germany (15, 17, 19, 22). At follow-up, sweat test decreased by a mean of 11 mmol/L (−38-+31) (p<0.0001, paired t-test), and most remained elevated above 60 mmol/L (**Supplemental Figure 7a**). Eight patients significantly decreased their sweat test by more than 20 mmol/L (**Supplemental Figure 7b**). Beta-adrenergic sweat production was not changed in 2 patients, but interestingly the response to cholinergic secretion was decreased (**Supplemental Figure 8**). pwCF improved their ppFEV_1_ by a mean of 17.5%, in the same range as that predicted from CFTR activity rescue, ranging between 1 and 59% (p<0.0001, paired t-test) (**Supplemental Figure 7c and 7d)**. This variability was not explained by the respiratory severity as there was no correlation with ppFEV_1_ at baseline nor the number of N1303K alleles (18(10.5)% change in heterozygotes *versus* 16.5% (14.4) in homozygotes). There was no correlation between sweat test variation and ppFEV_1_ change either.

Investigation of potential other genetic factors accounting for the variability in response to ETI was performed in non or transient responders from the French cohort (#6, #8, #9, #15, and #16) and in 2 patients from the Italian cohort (patient ME084 and ME087) respectively weak responder and best responder (**Supplemental Manuscript**). Only the frequent haplotype described in *cis* with N1303K (c.[744-33GATT[6];869+11C>T]) was observed (34) and no other *CFTR* variant was identified as a complex allele. Three of the five French patients carried variants in other genes: patient #6 was a compound heterozygote for *SERPINA1* Z and S variants, a genotype responsible for alpha1-antitrypsin deficiency ; Patient #15 carried a pathogenic variant in the *SLC26A4* gene, R185T, involved in Pendred syndrome (35), patient #16 carried a gain-of-function variant in the *SCNN1A* gene, R181W (36).

We succeeded to get approval for API in addition to ETI for patient (#3) who transiently improved ppFEV_1_ from 44% to 69% at 1 month ETI but subsequently progressively decreased to 62% of the normal at nine months ETI treatment. The patient was treated with API for two months, which was associated with a decrease in sputum production and an increase in ppFEV_1_ by 10%. The patient decided to stop after two months of treatment because of the taste of API and ppFEV_1_ dropped back down to its initial value.

## DISCUSSION

Our results confirm that N1303K can be corrected by ELX/TEZ. Importantly, they also point out that N1303K-CFTR and F508del-CFTR present distinct maturation defects and correction patterns. N1303K-CFTR is less prone to degradation and more distributed at the cell surface both in control and corrected conditions. Importantly, ELX/TEZ mainly induces a global stabilization of N1303K-CFTR without favoring its maturation and plasma membrane relocation, in contrast to F508del-CFTR. In the respiratory epithelium, N1303K-CFTR activity restoration by ETI is variable and can be increased with API, consistent with a strong gating defect and this is clinically relevant.

ETI has proved efficient for pwCF carrying N1303K and non-eligible mutations (14,15,17,18,19,20,21). The level of improvement was similar to that observed for F508del patients. The variability of the response was not explained by lung severity at baseline nor additional mutations in *cis* of N1303K (34). Interestingly, we identified S/Z alpha1 antitrypsin genotype and gain of function *SLC26A4* and *SCNN1A* (35, 36) variants in non or transient responders, suggesting that additional genes must be investigated in pwCF displaying an atypical response to ETI. As recently reported, importantly airway inflammation may also be a key factor in the rescue of N1303K-CFTR (37).

This contrasts with a faint impact on sweat chloride concentration, and no change in CFTR-dependent beta-adrenergic secretion, although ETI was active in a sweat gland derived cell line. The decrease in the fluid reabsorption in response to cholinergic stimulation suggests however a functional restoration in the sweat tubule.

Importantly, our results unravel that N1303K-CFTR and F508del-CFTR display different maturation defects. Indeed, at basal state, N1303K-CFTR showed greater band B amounts and was present at the cell surface as demonstrated by biochemical and functional approaches in different cell models. This was supported by the observation that in ∼1/3 patient HNECs, in control conditions, N1303K-CFTR activity was potentiated by IVA/API, as already reported by Phuan et al (24). Moreover, comparative N1303K *versus* F508del-CFTR interactomic studies suggested that uncorrected N1303K-CFTR interacts with proteins from the Golgi apparatus, endosomes and plasma membrane.

As reported by others (15, 16), we confirm that ELX-TEZ corrects N1303K-CFTR, and this is consistent with a gain in ppFEV_1_ as previously published (14). We also show that the pattern of correction is different from that of F508del-CFTR. First, ELX/TEZ increases the amount of surface N1303K-CFTR expression, but this is linked to a global increase of protein and not a specific impact on maturation or trafficking as neither the proportion of mature CFTR or total amount of protein at the surface were modified when normalized to total CFTR quantity. This response was in contrast with that of F508del-CFTR, whose correction enabled a clear switch to maturation associated with augmentation of the proportion of channels at the cell surface. Second, degradation rate of N1303K-CFTR was much slower than that of F508del-CFTR. Previous findings suggested that endoplasmic reticulum-associated degradation-resistant pools of N1303K-CFTR were found in ER-connected phagophores and were degraded through the autophagic pathway (22, 38). In our experimental conditions, the inhibition of the autophagic pathway did not enhance rescue by modulators, indicating that N1303K-CFTR engaged in the autophagic pathway is not rescuable.

At the functional level, increased N1303K-CFTR expression by ELX/TEZ was associated with a gain in CFTR channel activity but combining IVA with its co-potentiator API further increased CFTR activity by 5%. Importantly, in ∼ 20% of the N1303K-HNECs, CFTR activity was only improved after addition of API after IVA in corrected cells. These observations are clinically relevant as a rescue of 5% of CFTR activity *in vitro* is associated to 10% increase in ppFEV1 at short term (14), and indeed, API induced a ppFEV_1_ gain of 10% in a patient already treated with ETI. Apigenin is a plant-derived flavone, reported as a CFTR activator (39), considered for a therapeutic use since the 1950s for bronchodilation and now in humans as an adjuvant for cancer therapy (40).

## Conclusion

Our results suggest a different mechanism of action of ETI compared to F508del. These observations are particularly relevant to CF as they support CF theranostics bringing causal therapies to pwCF with rare mutations

## Supporting information

Supplemental manuscript

---

**Supplemental Figure 1.**
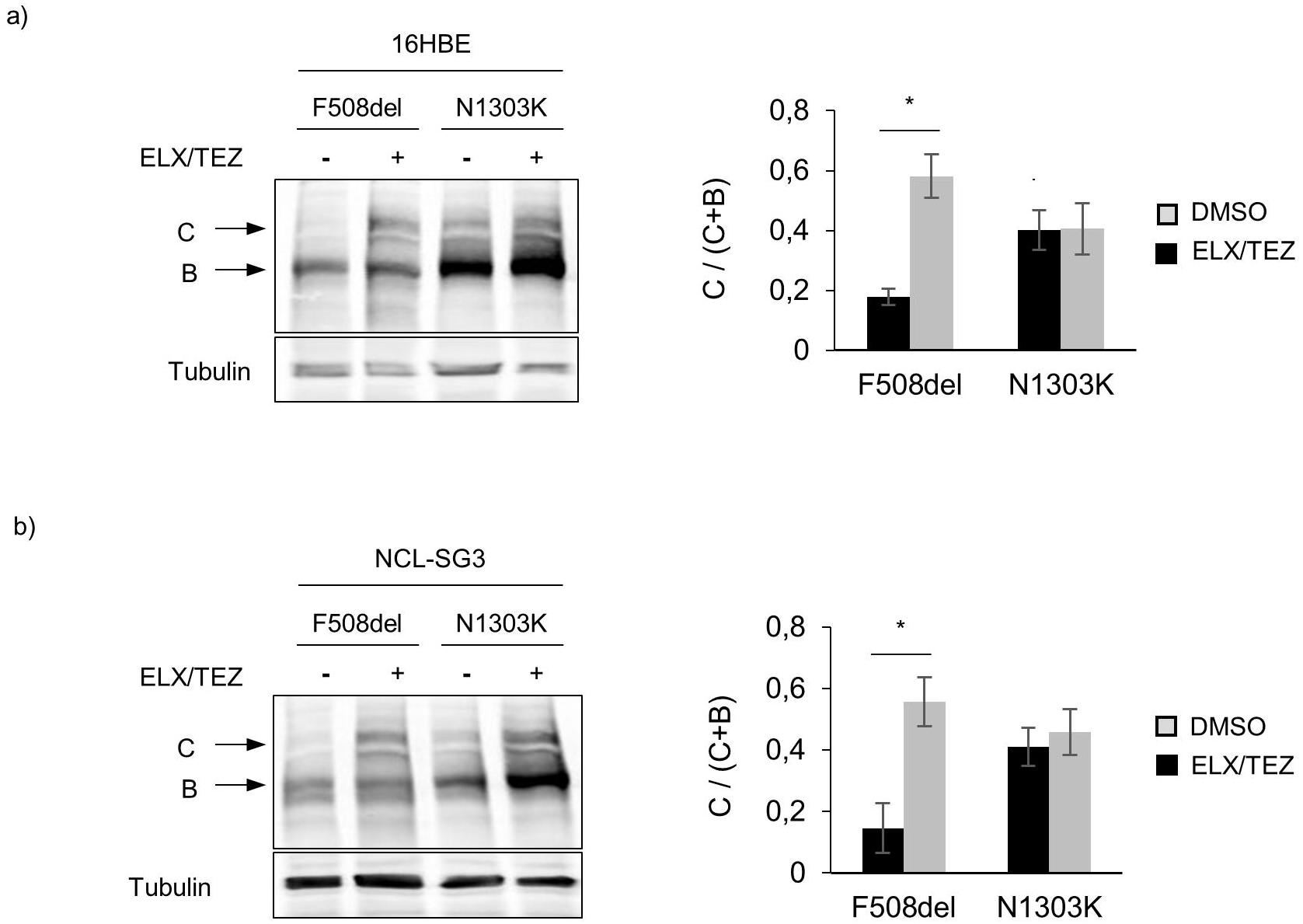
N1303K-CFTR correction in 16HBEge and NCL-SG3 cell lines. a) Representative Western blot images of N1303K-CFTR and F508del-CFTR from whole cell lysates from 16HBEge-G542X-CFTR cell lines transiently expressing N1303K-CFTR and incubated for 24 hours with DMSO (vehicle) or correctors elexacaftor/tezacaftor (ELX/TEZ at 3 µM each). b) Representative Western blot images of N1303K-CFTR and F508del-CFTR from whole cell lysates from NCL-SG3 cell lines transiently expressing N1303K-CFTR and incubated for 24 hours with DMSO (vehicle) or correctors (ELX/TEZ at 3 µM each). For a) and b) B band corresponds to the immature core glycosylated fraction and C to the mature, fully glycosylated fraction of the CFTR protein. Bar graphs correspond to the corresponding quantification of the C/(C + B) band ratio, expressed as mean ± standard deviation (SD) from a minimum of three independent experiments. Comparison by unpaired t-test, * for p<0.05.

**Supplemental Figure 2.**
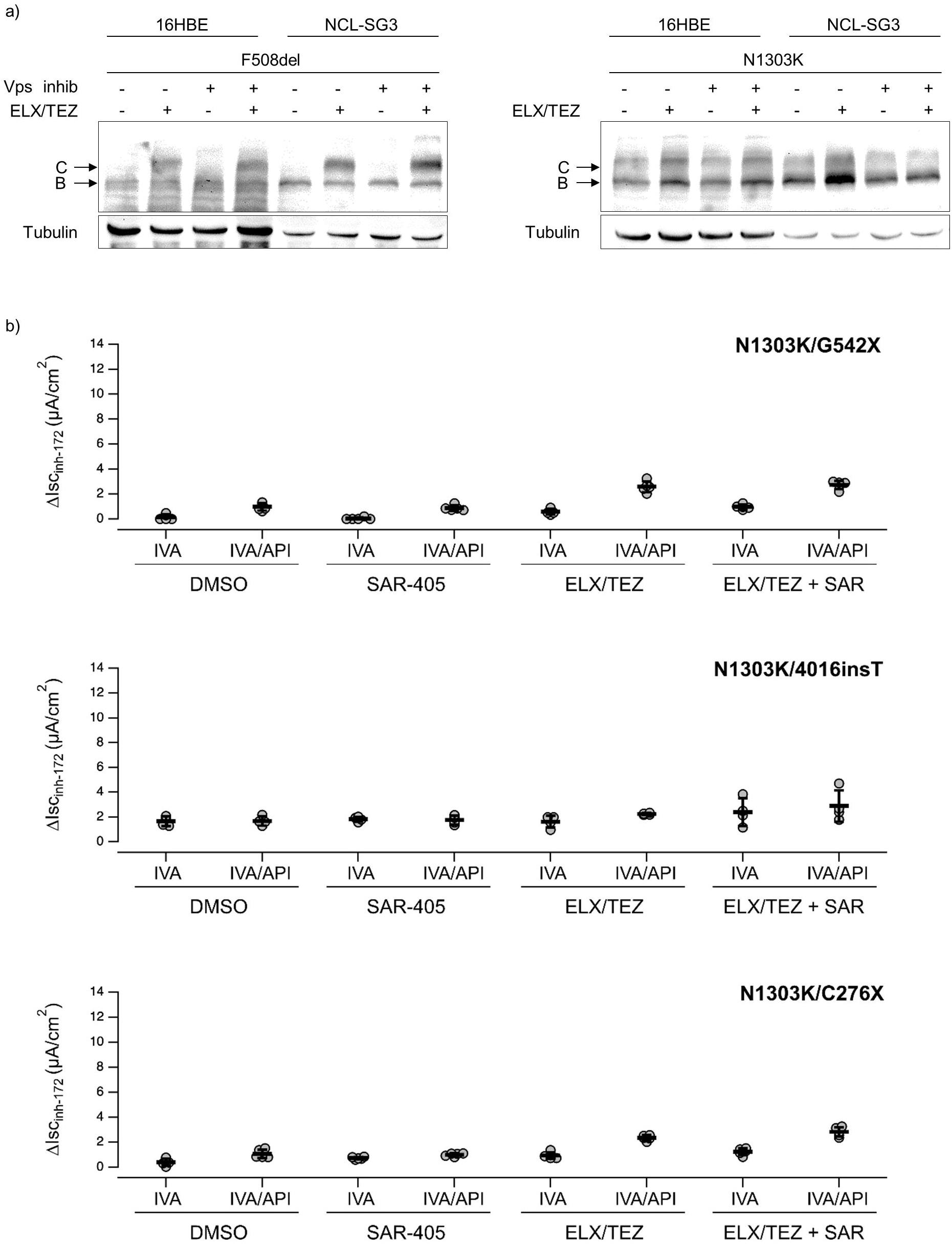
Impact of autophagy on N1303K-CFTR trafficking/maturation. a) Representative Western blot of 16HBEge-G542X-CFTR and NCL-SG3 cell lines transiently expressing F508del-CFTR (left panel) or N1303K-CFTR (right panel) and incubated for 24 hours with DMSO (vehicle) or correctors elexacaftor/tezacaftor (ELX/TEZ at 3 µM each) and/or with Vps inhibitor (10 µM) as indicated. B band corresponds to the immature core glycosylated fraction and C to the mature, fully glycosylated fraction of the CFTR protein. b) Scatter dot plot showing the summary of the results obtained during experiments performed with the short-circuit-current technique on nasal epithelial cells derived from 3 subjects compound heterozygous for N1303K and non-rescuable variant (G542X (donor ID: MI247), 4016insT (donor ID: MI248) and C276X (donor ID: MI259) as indicated). Prior to the experiments, epithelia were treated for 24 hrs with vehicle (DMSO), or the ELX/TEZ combination (3 µM/10 µM) in the absence or presence of the Vps inhibitor SAR-405 (2 µM, 3 hrs,). During the recordings, the epithelia were sequentially treated with amiloride (10 μM), CPT-cAMP (100 μM), ivacaftor (IVA, 1 μM), apigenin (API, 25 μM; where indicated), and the CFTR inhibitor-172 (inh-172; 20 μM). Data reported are the amplitude of the current blocked by inh-172 (ΔIsc_inh-172_). For each experimental condition the number of biological replicates was n = 4-6.

**Supplemental Figure 3.**
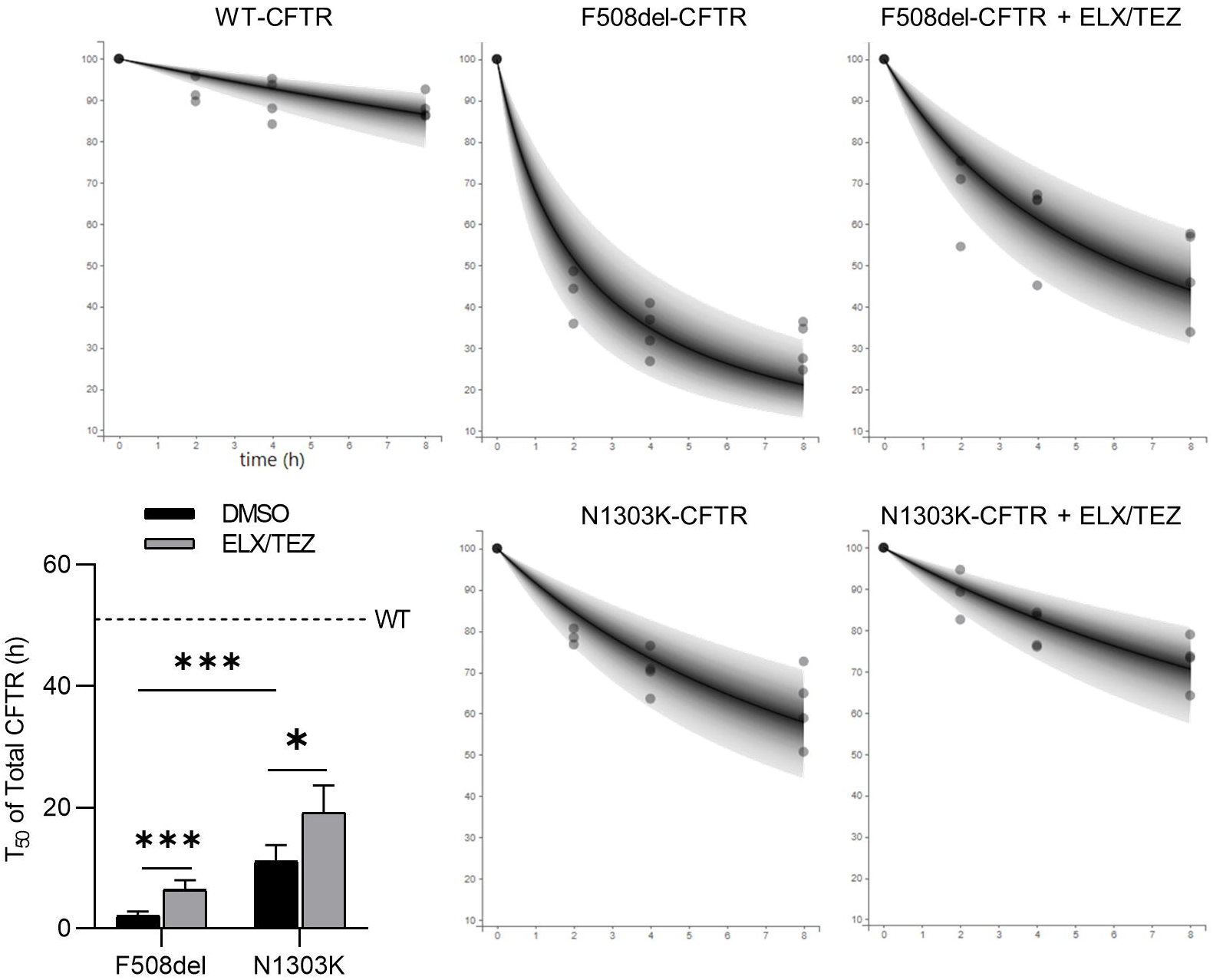
Modeling of total CFTR degradation from the Cycloheximide Chase Analysis. a) Prediction distribution plots derived from the modeling of total CFTR stability under cycloheximide (Norm/%T0) overlayed with the observed data (points). The areas represent the 95% prediction interval stratified according to experimental conditions. b) Estimated N1303K-CFTR and F508del-CFTR half-life according to different conditions. Data are presented as mean ± SEM. ** for p<0.01 and *** for p<0.001 (unpaired t-test, n=4).

**Supplemental Figure 4.**
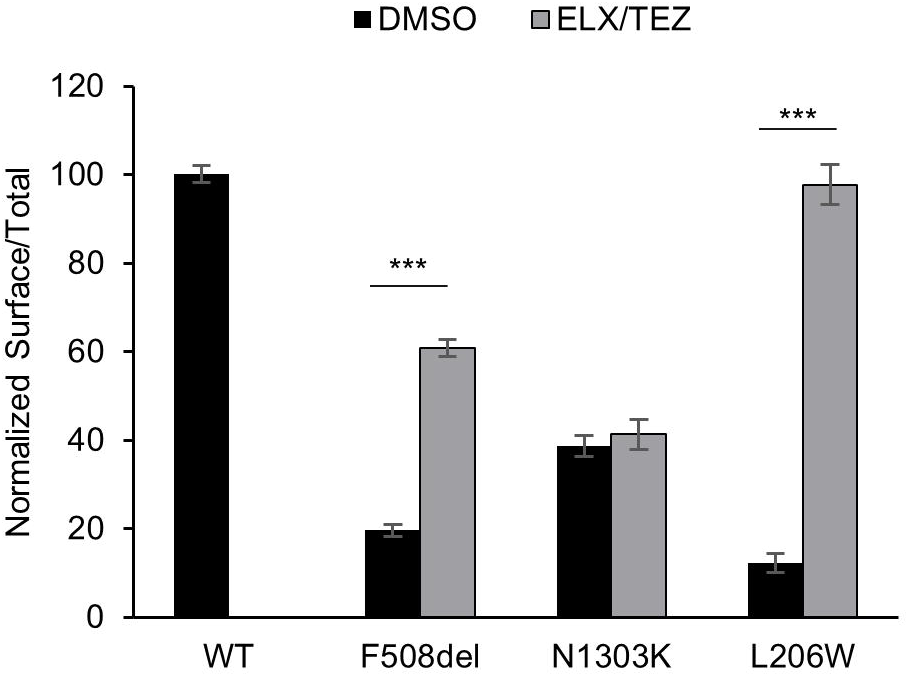
L206W-CFTR expression in response to ELX/TEZ. Nanoluciferase complementation assay quantification of Surface/Total ratio performed on HEK293 cells transfected with HiBiT-L206W-CFTR treated for 24 hours with either DMSO or correctors elexacaftor/tezacaftor (ELX/TEZ, 3 µM each) when indicated. Luminescent signals were normalized to HiBiT-WT-CFTR in DMSO conditions and data are presented as mean ± SEM. WT, F508del and N1303K shown for comparison. Stars indicate significant differences between the indicated comparisons of at least four independent experiments, unpaired t-test, ns for p> 0.05; *** for p<0.001.

**Supplemental Figure 5.**
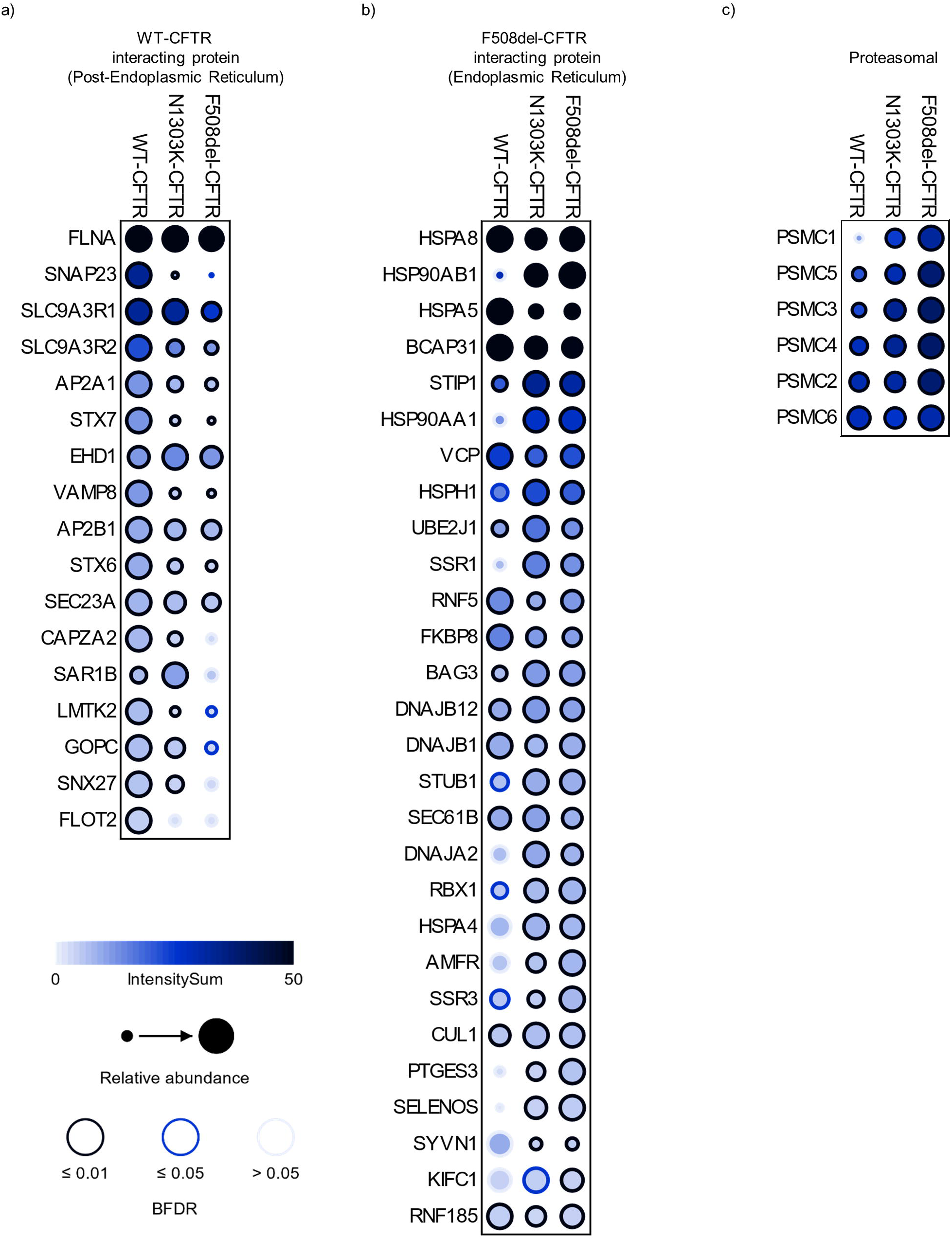
Analysis of proximal partners of N1303K-CFTR in comparison to WT or F508del-CFTR mutant using proximity labeling TurboID from HEK cells (n=3). List of CFTR interacting proteins were obtained from CyFi-map (https://cysticfibrosismap.github.io/). Data are represented as a dot plot: the color of the circle indicates the average intensity values, circle size displays the relative abundance of preys (interacting proteins) between WT and mutant CFTR, the colored edge of the circles indicates the BFDR or confidence of the interaction. Statistical analysis performed by SAINTexpress and visualization performed using ProHits-viz.

**Supplemental Figure 6.**
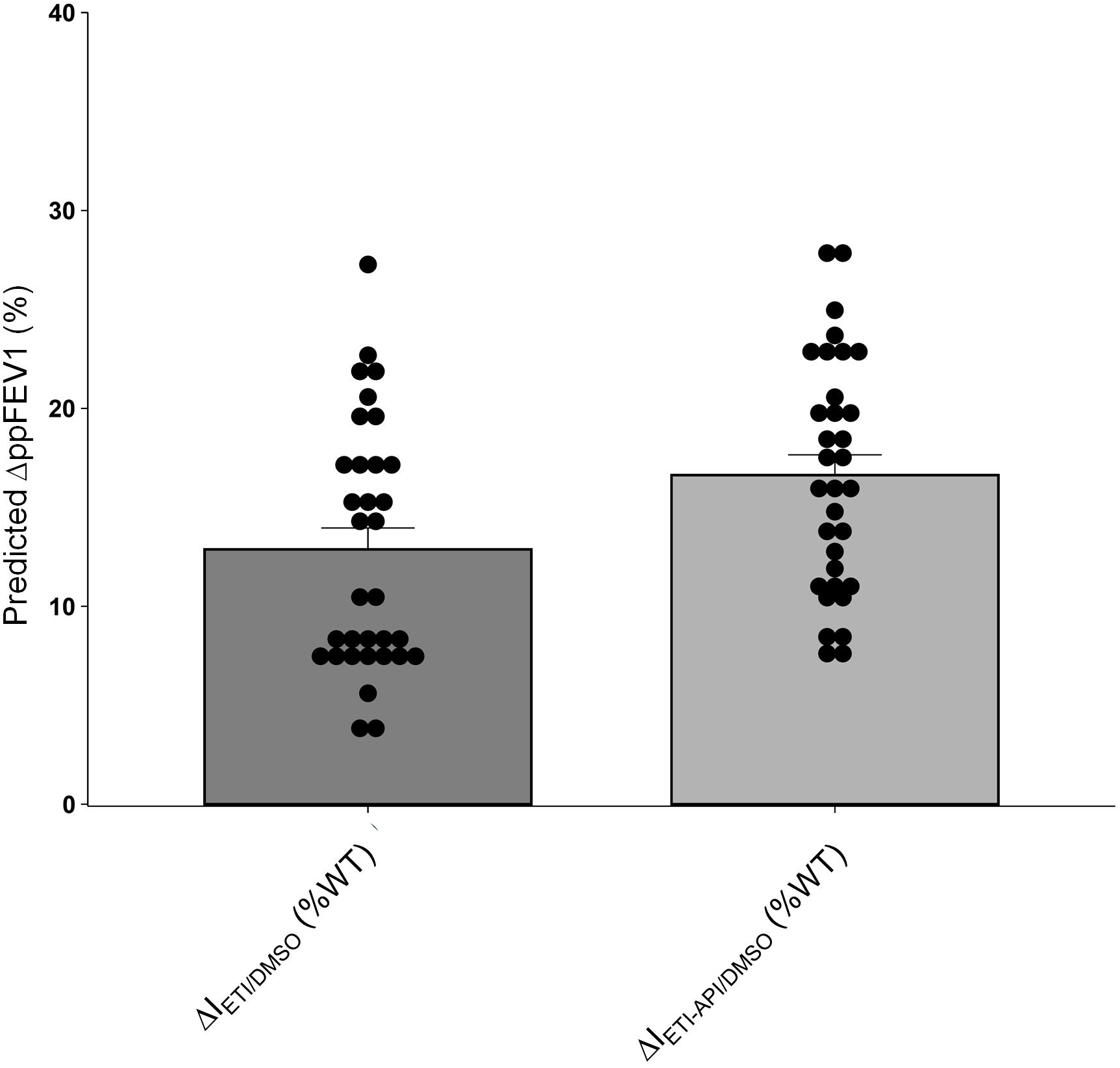
Predicted gain in Forced Expiratory Volume in pwCF carrying N1303K treated with ETI or ETI/API based on the level of CFTR activity rescue. Absolute gain in ppFEV_1_ expressed in percentage predicted _(_ΔppFEV_1_) is correlated to CFTR activity correction upon ETI, expressed in percentage of Wild Type (ΔI_ETI/DMSO_ (%WT)) where ΔI is Isc change after stimulation with cAMP agonists and its inhibition by Inh-172(ΔIsc_inh-172_) This is best described by following equation ΔppFEV1 = 30.9 × (1 − 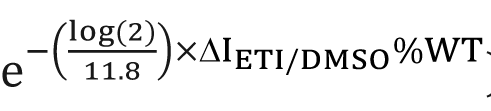) (14). Data are presented as mean± SEM with individual value for ETI (dark grey) and ETI/API combination (light grey).

**Supplemental Figure 7.**
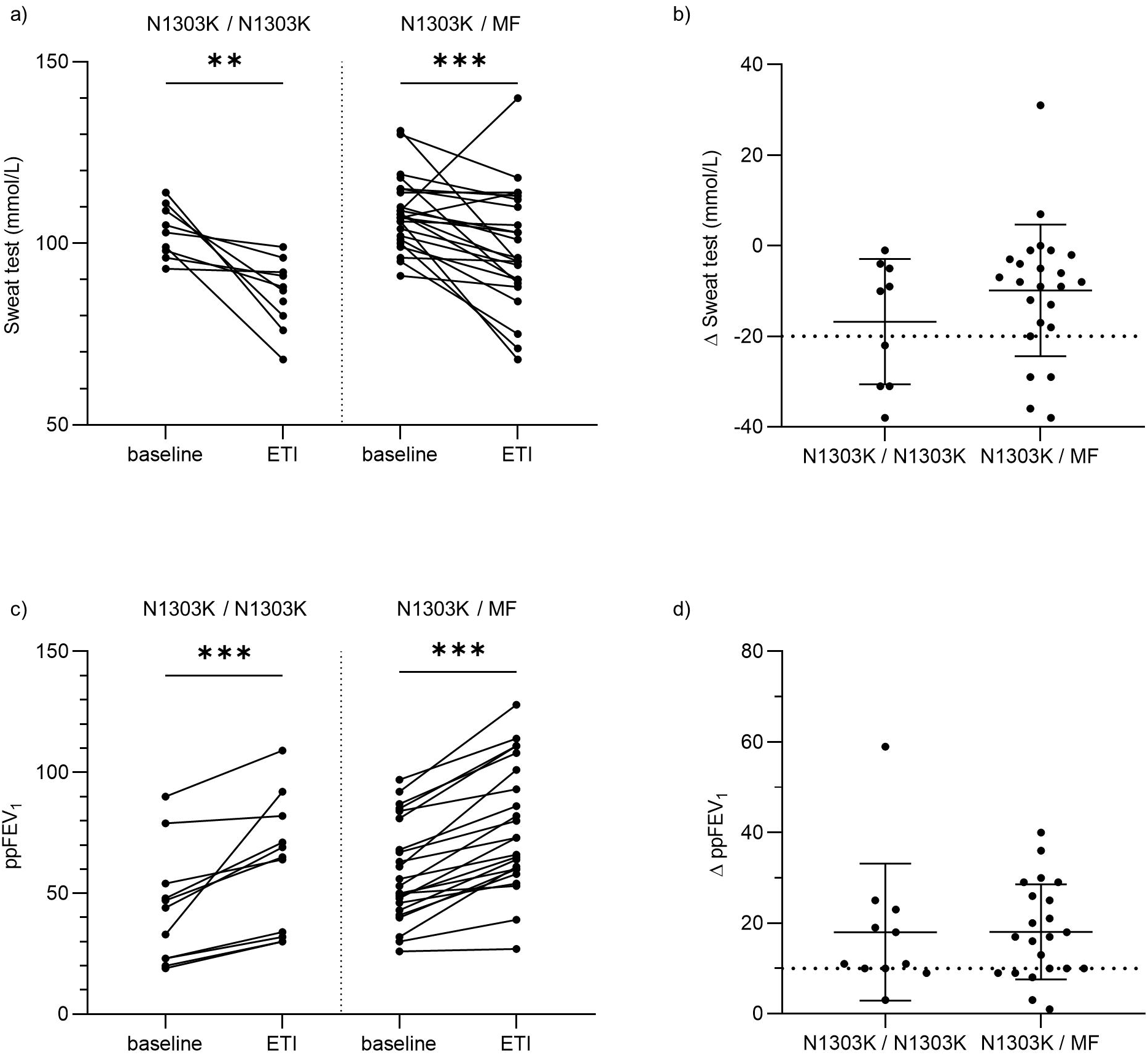
Change in sweat chloride concentration and percent predicted forced expiratory volume in 1 second in pwCF carrying the N1303K variant treated with ETI. Connected dot plots indicating a) the Sweat chloride concentration and b) the percent predicted forced expiratory volume in 1 second (ppFEV_1_) before treatment (Baseline) and under ETI (ETI) in homozygous N1303K patients (N1303K/N1303K) and patients carrying N1303K in *trans* of a minimal function variant (N1303K/MF). Scatter dot plots indicate the corresponding c) Sweat test variation (ΔSweat test) and d) variation of ppFEV_1_ (ΔppFEV_1_) under ETI. a) and c) Comparison by Wilcoxon matched-pairs signed rank test, b) and d) comparison by unpaired t test. ** for p<0.01 and *** for p<0.001.

**Supplemental Figure 8.**
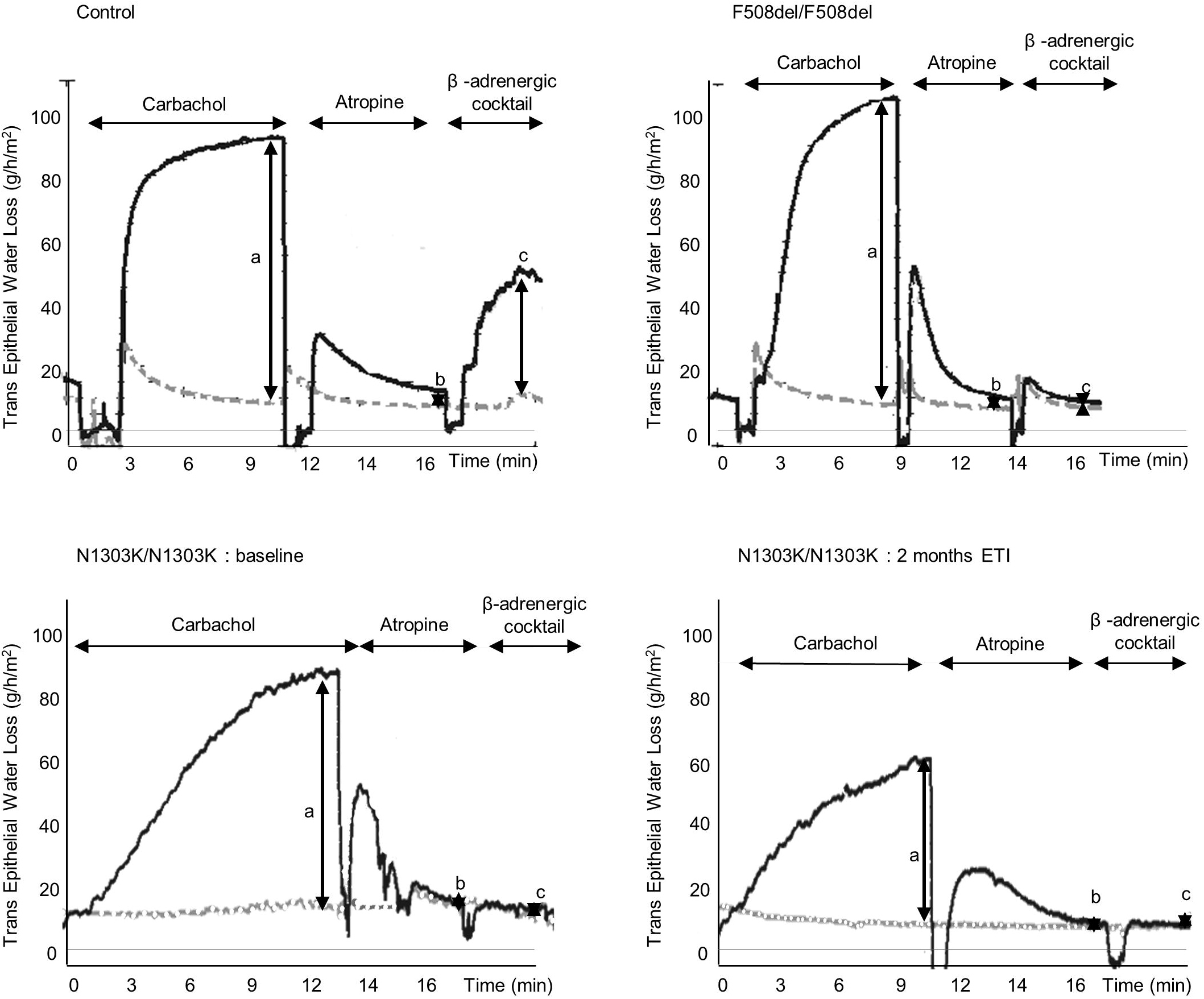
Beta-adrenergic sweat secretion in a healthy subject, a pwCF homozygous for F508del and a N1303K homozygous pwCF before initiation of ETI and at one-month ETI treatment. Transepithelial Water loss (TEWL) is measured as kg/m^2^/h on the Y-axis over time in minutes on the X-axis. Measurement of baseline water loss is followed by abrupt movement artifacts associated with lifting the evapometry probe to inject carbachol (0.1 ml = 0.01 mg) intradermally. After maximal response (carbachol peak in g/m^2^/h), atropine (0.2 ml = 8.8 mg) is injected to block the cholinergic sweat secretion. After baseline stabilization, the beta-adrenergic cocktail comprising atropine 8.8 mg, isoproterenol 4.4 mg, and aminophylline 0.93mg is injected. The maximal evaporative beta-adrenergic-response is reported as the mean secretion (beta-adrenergic peak in kg/m^2^/h) and the ratio to the carbachol peak. Representative trace of carbachol peak, beta-adrenergic peak, ratio carbachol/beta-adrenergic in a) a control subject (Control) Carbachol peak: 75 g/m^2^/h. Beta-adrenergic peak: 52 g/m^2^/h. Ratio beta-adrenergic/carbachol: 0.69 a) F508del homozygous patient (F508del/F508del) Carbachol peak: 85.7 g/m^2^/h. Beta-adrenergic peak: 0 kg/m^2^/h. Ratio beta-adrenergic/carbachol: 0 and c) and d) a N1303K homozygous patient (N1303K/N1303K) a) before treatment Carbachol peak: 76.9 kg/m^2^/h. Beta-adrenergic peak: 2.9 kg/m^2^/h. Ratio beta-adrenergic/carbachol: 0.04 a) 1 month treatment with ETI. Carbachol peak: 44.4 kg/m^2^/h. Beta-adrenergic peak: 0 kg/m^2^/h. Ratio beta-adrenergic/ carbachol: 0.02

## Notes

### Competing Interest Statement

Stefano Pantano declares Vertex pharmaceuticals support in sample collection without financial contribution.
Stefano Costa declares payment or honoraria for speakers bureaus from Vertex pharmaceuticals.
Sonia Volpi declares payment of honoraria for lectures, presentations, speakers bureaus, manuscript writing or educational events from Vertex pharmaceuticals and DMF Pharma FoodAR and support for attending meetings and/or travel from Chiesi. Stephanie Bui declares participation in the protocoles of Vertex pharmaceuticals studies as principal investigator.
Clemence Martin reports payment or honoraria for lectures, presentations, speakers bureaus, manuscript writing or educational events from Chiesi, Astra Zeneca, Boehringer Ingelheim, GSK. Support for attending meetings and/or travel from Chiesi, Boehringer Ingelheim.
Nicoletta Pedemonte declares payment or honoraria from Vertex Pharmaceuticals for lectures, presentations, speakers bureaus, manuscript writing or educational events - Speaker for a lecture at the 45th European Cystic Fibrosis Society Conference, Rotterdam, June 2022.
Pierre Regis Burgel reports grants from Vertex pharmaceuticals, GSK, outside the submitted work.
Luis J.V. Galietta declares Patents planned, issued or pending. Compounds described are not present in the submitted paper.
Isabelle Sermet-Gaudelus reports support for the present manuscript from Vaincre La Mucoviscidose and Mucoviscidose ABCF2. Isabelle Sermet-Gaudelus also reports, outside the submitted work, grants from Agence Nationale pour la Recherche, Assistance Publique Hopitaux de Paris, Vertex Innovation Award; consulting fees and travel support from Vertex therapeutics.

## Bibliography

1. Graeber SY, Mall MA. The future of cystic fibrosis treatment: from disease mechanisms to novel therapeutic approaches. Lancet. 2023 Sep 30;402(10408):1185–1198. doi: 10.1016/S0140-6736(23)01608-2. Epub 2023 Sep 9. PMID: 37699417.

2. Cutting, G. Cystic fibrosis genetics: from molecular understanding to clinical application. Nat Rev Genet 16, 45–56 (2015).

3. Pranke I, Golec A, Hinzpeter A, Edelman A, Sermet-Gaudelus I. Emerging Therapeutic Approaches for Cystic Fibrosis. From Gene Editing to Personalized Medicine. Front Pharmacol. 2019 Feb 27;10:121. doi: 10.3389/fphar.2019.00121. PMID: 30873022; PMCID: PMC6400831

4. Middleton PG, Mall MA, Dřevínek P, Lands LC, McKone EF, Polineni D, Ramsey BW, Taylor-Cousar JL, Tullis E, Vermeulen F, Marigowda G, McKee CM, Moskowitz SM, Nair N, Savage J, Simard C, Tian S, Waltz D, Xuan F, Rowe SM, Jain R; VX17-445-102 Study Group. Elexacaftor-Tezacaftor-Ivacaftor for Cystic Fibrosis with a Single Phe508del Allele. N Engl J Med. 2019 Nov 7;381(19):1809–1819. doi: 10.1056/NEJMoa1908639. Epub 2019 Oct 31. PMID: 31697873; PMCID: PMC7282384.

5. US Food and Drug Administration. Trikafta Label. Available online: https://www.accessdata.fda.gov/drugsatfda_docs/label/2023/217660s000lbl.pdf (accessed on 16 July 2023).

6. Prontera P, Isidori I, Mencarini V, Pennoni G, Mencarelli A, Stangoni G, Di Cara G, Verrotti A. A Clinical and Molecular Survey of 62 Cystic Fibrosis Patients from Umbria (Central Italy) Disclosing a High Frequency (2.4%) of the 2184insA Allele: Implications for Screening. Public Health Genomics. 2016;19(6):336–341. doi: 10.1159/000450849. Epub 2016 Oct 12. PMID: 27728908.

7. Federici S, Iron A, Reboul MP, Desgeorges M, Claustres M, Bremont F, Bieth E. Etude du gène CFTR chez 207 patients du Sud-Ouest de la France atteints de mucoviscidose: fréquence élevée des mutations N1303K et 1811 + 1,6 kbA > G [CFTR gene analyis in 207 patients with cystic fibrosis in southwest France: high frequency of N1303K and 1811+1.6bA>G mutations]. Arch Pediatr. 2001 Feb;8(2):150–7. French. doi: 10.1016/s0929-693x(00)00177-9. PMID: 11232455.

8. Noel S, Sermet-Gaudelus I, Sheppard DN. N1303K: Leaving no stone unturned in the search for transformational therapeutics. Journal of Cystic Fibrosis 2018: 17(5): 555-557

9. Veit G, Roldan A, Hancock MA, Da Fonte DF, Xu H, Hussein M, Frenkiel S, Matouk E, Velkov T, Lukacs GL. Allosteric folding correction of F508del and rare CFTR mutants by elexacaftor-tezacaftorivacaftor (Trikafta) combination. JCI Insight 2020: 5(18)

10. Destefano S, Gees M, Hwang T-C. Physiological and pharmacological characterization of the N1303K mutant CFTR. Journal of Cystic Fibrosis 2018: 17(5): 573–581.

11. Bihler H, Sivachenko A, Millen L, Bhatt P, Patel AT, Chin J, et al. In Vitro Modulator Responsiveness of 655 CFTR Variants Found in People With CF. bioRxiv. 2023:2023.07.07.548159.

12. Durmowicz AG, Lim R, Rogers H, Rosebraugh CJ, Chowdhury BA. The U.S. Food and Drug Administration’s Experience with Ivacaftor in Cystic Fibrosis. Establishing Efficacy Using In Vitro Data in Lieu of a Clinical Trial. Annals of the American Thoracic Society. 2017;15(1):1–2.

13. Laselva O, Bartlett C, Gunawardena TNA, Ouyang H, Eckford PDW, Moraes TJ, Bear CE, Gonska T. Rescue of multiple class II CFTR mutations by elexacaftor+tezacaftor+ivacaftor mediated in part by the dual activities of elexacaftor as both corrector and potentiator. European Respiratory Journal 2021: 57(6): 2002774.

14. Dreano E, Burgel PR, Hatton A, Bouazza N, Chevalier B, Macey J, Leroy S, Durieu I, Weiss L, Grenet D, Stremler N, Ohlmann C, Reix P, Porzio M, Roux Claude P, Rémus N, Douvry B, Montcouquiol S, Cosson L, Mankikian J, Languepin J, Houdouin V, Le Clainche L, Guillaumot A, Pouradier D, Tissot A, Priou P, Mély L, Chedevergne F, Lebourgeois M, Lebihan J, Martin C, Zavala F, Da Silva J, Lemonnier L, Kelly-Aubert M, Golec A, Foucaud P, Marguet C, Edelman A, Hinzpeter A, di Carli P, Girodon E, Sermet-Gaudelus I, Pranke I. Theratyping Cystic Fibrosis patients to guide Elexacaftor-Tezacaftor-Ivacaftor out of label prescription. Eur Respir J. 2023 Sep 11:2300110. doi: 10.1183/13993003.00110-2023. Epub ahead of print. PMID: 37696564.

15. Huang Y, Paul G, Lee J, Yarlagadda S, McCoy K, Naren AP. Elexacaftor/Tezacaftor/Ivacaftor Improved Clinical Outcomes in a Patient with N1303K-CFTR Based on *In Vitro* Experimental Evidence. Am J RespirCrit Care Med. 2021 Nov 15;204(10):1231–1235. doi: 10.1164/rccm.202101-0090LE. PMID: 34379998; PMCID: PMC8759307.

16. Ensinck MM, De Keersmaecker L, Ramalho AS, Cuyx S, Van Biervliet S, Dupont L, Christ F, Debyser Z, Vermeulen F, Carlon MS. Novel CFTR modulator combinations maximise rescue of G85E and N1303K in rectal organoids. ERJ Open Res. 2022 Apr 19;8(2):00716–2021. doi: 10.1183/23120541.00716-2021. PMID: 35449760; PMCID: PMC9016267.

17. Sadras I, Kerem E, Livnat G, Sarouk I, Breuer O, Reiter J, Gileles-Hillel A, Inbar O, Cohen M, Gamliel A, Stanleigh N, Gunawardena T, Bartlett C, Gonska T, Moraes T, Eckford PDW, Bear CE, Ratjen F, Kerem B, Wilschanski M, Shteinberg M, Cohen-Cymberknoh M. Clinical and functional efficacy of elexacaftor/tezacaftor/ivacaftor in people with cystic fibrosis carrying the N1303K mutation. J Cyst Fibros. 2023 Jun 16:S1569–1993(23)00178-9. doi: 10.1016/j.jcf.2023.06.001. Epub ahead of print. PMID: 37331863

18. Burgel PR, Sermet-Gaudelus I, Durieu I, Kanaan R, Macey J, Grenet D, Porzio M, Coolen-Allou N, Chiron R, Marguet C, Douvry B, Dufeu N, Danner-Boucher I, Foucaud P, Lemonnier L, Girodon E, Da Silva J, Martin C; French CF Reference Network study group. The French Compassionate Program of elexacaftor-tezacaftor-ivacaftor in people with cystic fibrosis with advanced lung disease and no F508del CFTR variant. Eur Respir J. 2023 Feb 16:2202437. doi: 10.1183/13993003.02437-2022. Epub ahead of print. PMID: 36796836.

19. Graeber SY, Balázs A, Ziegahn N, Rubil T, Vitzthum C, Piehler L, Drescher M, Seidel K, Rohrbach A, Röhmel J, Thee S, Duerr J, Mall MA, Stahl M. Personalized CFTR Modulator Therapy for *G85E* and *N1303K* Homozygous Patients with Cystic Fibrosis. Int J Mol Sci. 2023 Aug 2;24(15):12365. doi: 10.3390/ijms241512365. PMID: 37569738; PMCID: PMC10418744.

20. https://news.vrtx.com/news-releases/news-release-details/vertex-announces-european-medicines-agency-validation-marketing, accessed 24/11/2023

21. Burgel PR, Sermet-Gaudelus I, Girodon E, Kanaan R, Le Bihan J, Remus N, Ravoninjatovo B, Grenet D, Porzio M, Houdouin V, Le Clainche-Viala L, Durieu I, Nove-Josserand R, Languepin J, Coltey B, Guillaumot A, Audousset C, Chiron R, Weiss L, Fajac I, Da Silva J, Martin C; French CF Reference Network study group. Gathering real-world compassionate data to expand eligibility to elexacaftor-tezacaftor-ivacaftor in people with cystic fibrosis with N1303K or other rare CFTR variants: a viewpoint. Eur Respir J. 2024 Jan 19:2301959. doi: 10.1183/13993003.01959-2023. Epub ahead of print. PMID: 38242629.

22. Liu Q, Sabirzhanova I, Yanda MK, Bergbower EAS, Boinot C, Guggino WB, Cebotaru L. Rescue of CFTR NBD2 mutants N1303K and S1235R is influenced by the functioning of the autophagosome. Journal of Cystic Fibrosis 2018: 17(5): 582–594. 26.

23. Liu F, Zhang Z, Csanády L, Gadsby DC, Chen J. Molecular Structure of the Human CFTR Ion Channel. Cell. 2017 Mar 23;169(1):85–95.e8. doi: 10.1016/j.cell.2017.02.024. PMID: 28340353.

24. Phuan P-W, Son J-H, Tan J-A, Li C, Musante I, Zlock L, Nielson DW, Finkbeiner WE, Kurth MJ, Galietta LJ, Haggie PM, Verkman AS. Combination potentiator (‘co-potentiator’) therapy for CF caused by CFTR mutants, including N1303K, that are poorly responsive to single potentiators. Journal of Cystic Fibrosis 2018: 17(5): 595–606

25. Phuan PW, Tan JA, Rivera AA, Zlock L, Nielson DW, Finkbeiner WE, Haggie PM, Verkman AS. Nanomolar-potency ’co-potentiator’ therapy for cystic fibrosis caused by a defined subset of minimal function CFTR mutants. Sci Rep. 2019 Nov 27;9(1):17640. doi: 10.1038/s41598-019-54158-2. PMID: 31776420; PMCID: PMC6881293.

26. Valley, H. C. et al. Isogenic cell models of cystic fibrosis-causing variants in natively expressing pulmonary epithelial cells. J. Cyst. Fibros. 18, 476–483 (2019).

27. Servetnyk Z, Roomans GM. Chloride transport in NCL-SG3 sweat gland cells: channels involved. Exp Mol Pathol. 2007 Aug;83(1):47–53. doi: 10.1016/j.yexmp.2007.02.003. Epub 2007 Feb 27. PMID: 17383636

28. Sondo E, Cresta F, Pastorino C, Tomati V, Capurro V, Pesce E, Lena M, Iacomino M, Baffico AM, Coviello D, Bandiera T, Zara F, Galietta LJV, Bocciardi R, Castellani C, Pedemonte N. The L467F-F508del Complex Allele Hampers Pharmacological Rescue of Mutant CFTR by Elexacaftor/Tezacaftor/Ivacaftor in Cystic Fibrosis Patients: The Value of the Ex Vivo Nasal Epithelial Model to Address Non-Responders to CFTR-Modulating Drugs. Int J Mol Sci. 2022 Mar 15;23(6):3175. doi: 10.3390/ijms23063175. PMID: 35328596; PMCID: PMC8952007.

29. Tomati V, Costa S, Capurro V, Pesce E, Pastorino C, Lena M, Sondo E, Di Duca M, Cresta F, Cristadoro S, Zara F, Galietta LJV, Bocciardi R, Castellani C, Lucanto MC, Pedemonte N. Rescue by elexacaftor-tezacaftor-ivacaftor of the G1244E cystic fibrosis mutation’s stability and gating defects are dependent on cell background. J Cyst Fibros. 2023 May;22(3):525–537. doi: 10.1016/j.jcf.2022.12.005. Epub 2022 Dec 19. PMID: 36543707.

30. Terlizzi V, Pesce E, Capurro V, Tomati V, Lena M, Pastorino C, Bocciardi R, Zara F, Centrone C, Taccetti G, Castellani C, Pedemonte N. Clinical Consequences and Functional Impact of the Rare S737F CFTR Variant and Its Responsiveness to CFTR Modulators. Int J Mol Sci. 2023 Mar 31;24(7):6576. doi: 10.3390/ijms24076576. PMID: 37047546; PMCID: PMC10095403.

31. Dixon, A. S. et al. NanoLuc complementation reporter optimized for accurate measurement of protein interactions in cells. ACS Chem. Biol. 11, 400–408 (2016).

32. Chevalier B, Baatallah N, Najm M, Castanier S, Jung V, Pranke I, Golec A, Stoven V, Marullo S, Antigny F, Guerrera IC, Sermet-Gaudelus I, Edelman A, Hinzpeter A. Differential CFTR-Interactome Proximity Labeling Procedures Identify Enrichment in Multiple SLC Transporters. Int J Mol Sci. 2022 Aug 11;23(16):8937. doi: 10.3390/ijms23168937. PMID: 36012204; PMCID: PMC9408702.

33. Nguyen-Khoa T, Hatton A, Drummond D, Aoust L, Schlatter J, Martin C, Ramel S, Kiefer S, Gachelin E, Stremler N, Cosson L, Gabsi A, Remus N, Benhamida M, Hadchouel A, Fajac I, Munck A, Girodon E, Sermet-Gaudelus I. Reclassifying inconclusive diagnosis for cystic fibrosis with new generation sweat test. Eur Respir J. 2022 Aug 4;60(2):2200209. doi: 10.1183/13993003.00209-2022. PMID: 35777769.

34. Farhat R, El-Seedy A, Pasquet MC, Corbani S, Megarbané A, Kitzis A, Ladeveze V. Three Complex alleles associated with N1303K mutation and their molecular consequences. Cell Mol Biol (Noisy-le-grand). 2022 Apr 30;68(4):52–59. doi: 10.14715/cmb/2022.68.4.7. PMID: 35988290.

35. Chattaraj P, Reimold FR, Muskett JA, Shmukler BE, Chien WW, Madeo AC, Pryor SP, Zalewski CK, Butman JA, Brewer CC, Kenna MA, Alper SL, Griffith AJ. Use of SLC26A4 mutation testing for unilateral enlargement of the vestibular aqueduct. JAMA Otolaryngol Head Neck Surg. 2013 Sep;139(9):907–13. doi: 10.1001/jamaoto.2013.4185. Erratum in: JAMA Otolaryngol Head Neck Surg. 2014 Dec;140(12):1212. PMID: 24051746.

36. Azad AK, Rauh R, Vermeulen F, Jaspers M, Korbmacher J, Boissier B, Bassinet L, Fichou Y, des Georges M, Stanke F, De Boeck K, Dupont L, Balascáková M, Hjelte L, Lebecque P, Radojkovic D, Castellani C, Schwartz M, Stuhrmann M, Schwarz M, Skalicka V, de Monestrol I, Girodon E, Férec C, Claustres M, Tümmler B, Cassiman JJ, Korbmacher C, Cuppens H. Mutations in the amiloride-sensitive epithelial sodium channel in patients with cystic fibrosis-like disease. Hum Mutat. 2009 Jul;30(7):1093–103. doi: 10.1002/humu.21011. PMID: 19462466.

37. Gentzsch M, Baker B, Cholon DM, Kam CW, McKinzie CJ, Despotes KA, Boyles SE, Quinney NL, Esther CR Jr, Ribeiro CMP. Cystic fibrosis airway inflammation enables elexacaftor/tezacaftor/ivacaftor-mediated rescue of N1303K *CFTR* mutation. ERJ Open Res. 2024 Jan 15;10(1):00746–2023. doi: 10.1183/23120541.00746-2023. PMID: 38226069; PMCID: PMC10789252

38. He L, Kennedy AS, Houck S, Aleksandrov A, Quinney NL, Cyr-Scully A, Cholon DM, Gentzsch M, Randell SH, Ren HY, Cyr DM. DNAJB12 and Hsp70 triage arrested intermediates of N1303K-CFTR for endoplasmic reticulum-associated autophagy. Mol Biol Cell. 2021 Apr 1;32(7):538–553. doi: 10.1091/mbc.E20-11-0688. Epub 2021 Feb 3. PMID: 33534640; PMCID: PMC8101465..

39. Illek B, Lizarzaburu ME, Lee V, Nantz MH, Kurth MJ, Fischer H. Structural determinants for activation and block of CFTR-mediated chloride currents by apigenin. Am J Physiol Cell Physiol. 2000 Dec;279(6):C1838–46. doi: 10.1152/ajpcell.2000.279.6.C1838. PMID: 11078699.

40. DeRango-Adem EF, Blay J. Does Oral Apigenin Have Real Potential for a Therapeutic Effect in the Context of Human Gastrointestinal and Other Cancers? Front Pharmacol. 2021 May 18;12:681477. doi: 10.3389/fphar.2021.681477. PMID: 34084146; PMCID: PMC8167032.

