## Supplemental manuscript for "Beyond Kaftrio : mechanistic insights to maximize N1303K-CFTR rescue"

**# co-last**

****co-corresponding authors**

**Corresponding authors**:

Isabelle Sermet-Gaudelus ; Inserm U1151 - CNRS UMR 8253 – team 2- Faculté de Médecine Paris Descartes – 160 rue Vaugirard, Paris 75015, France ;; Tel: 0033 172606409

Nicoletta Pedemonte : UOC Genetica Medica, IRCCS Istituto Giannina Gaslini - Dirigente Biologo, Via Gerolamo Gaslini 5, 16147 Genova, Italy ;; Tel : +39 010 5636 3178

[1] INSERM, CNRS, Institut Necker Enfants Malades, Paris, France
[2] Université Paris-Cité, Paris, France
[3] UOC Genetica Medica, IRCCS Istituto Giannina Gaslini, Genova, Italy
[4] URC/CIC Paris Descartes Necker Cochin and Hôpital Tarnier, Paris, France
[5] Department of Neurosciences, Rehabilitation, Ophthalmology, Genetics, Maternal and Child Health (DINOGMI), University of Genoa, Genova, Italy
[6] UOSD CRR Fibrosi Cistica, P.O. San Liberatore, Atri, Italy
[7] Department of Pediatric Medicine, Meyer Children's Hospital IRCCS, Cystic Fibrosis Regional Reference Center, Florence, Italy
[8] Centro Hub Fibrosi Cistica, Azienda Ospedaliera Universitaria Policlinico G. Martino, Messina, Italy
[9] University Hospital "G Martino", Department of Pediatrics, Messina, Italy
[10] Department of Pediatrics, Cystic Fibrosis Center, Fondazione IRCCS Ca' Granda, Ospedale Maggiore Policlinico, Milan, Italy

[11] Department of Pediatrics, Cystic Fibrosis Regional Support Center, University of Brescia, ASST Spedali Civili Brescia, Brescia, Italy
[12] Institute for Maternal and Child Health-IRCCS "Burlo Garofolo", Trieste, Italy
[13] Cystic Fibrosis Regional Centre, Unit of Emerging and Immunosuppressed Infectious Diseases, Department of Gastroenterology and Transplantation, Azienda, Italy
[14] Ospedaliero-Universitaria ‘Ospedali Riuniti’, Ancona, Italy
[15] Cystic Fibrosis Center of Verona, Azienda Ospedaliera Universitaria Integrata, Verona, Italy
[16] Necker Proteomics, Université Paris Cité - Structure Fédérative de Recherche Necker, INSERM US24/CNRS UAR3633, Paris, France
[17] Cystic Fibrosis National Pediatric Reference Center, Pneumo-Allergologie Pédiatrique, Hôpital Necker Enfants Malades, AP-HP, Paris, France
[18] Centre de Ressources et de Compétence de la Mucoviscidose Enfants, Hôpital de Clocheville, Tours, France
[19] Centre de Ressources et de Compétence de la Mucoviscidose, CHU Pellegrin, Bordeaux, France
[20] Centre de Ressources et de Compétence de la Mucoviscidose Adulte, Centre de Perharidy, Roscoff, France
[21] Centre de Ressources et de Compétence de la Mucoviscidose Pédiatrique, CHU, Strasbourg, France
[22] Centre de Ressources et de Compétence de la Mucoviscidose, Hôpital Foch, Suresnes, France
[23] Centre de Ressources et de Compétence de la Mucoviscidose Pédiatrique, Hôpital Robert Debré, Paris, France
[24] Centre de Ressources et de Compétence de la Mucoviscidose Mixte, CHIC, Créteil, France
[25] Centre de Ressources et de Compétence de la Mucoviscidose, American Memoral Hospital, Reims, France
[26] Centre de Ressources et de Compétence de la Mucoviscidose, Institut Cœur Poumons, Lille, France
[27] CHU de Nancy - Hôpitaux de Brabois, Nancy, France
[28] Centre de Ressources et de Compétence de la Mucoviscidose Adulte, Centre Hospitalier Jean Minjoz, Besancon, France
[29] Centre de Ressources et de Compétence de la Mucoviscidose Enfants, Hôpital d'Enfants de la Timone, Marseille, France
[30] Centre de Ressources et de Compétence de la Mucoviscidose Enfants, Hôpital Trousseau, Paris, France
[31] Centre de Ressources et de Compétence de la Mucoviscidose, CHU Estaing, Clermont-Ferrand, France
[32] Centre de ressources et de compétences pour la mucoviscidose, Hôpital des enfants, CHU Toulouse, Toulouse, France
[33] Centre de Ressources et de Compétence de la Mucoviscidose Pédiatrique, Hospices Civils de Lyon, Bron, France
[34] Centre de Référence Adulte de la Mucoviscidose, Hospices Civils de Lyon, Université de Lyon, Lyon, France
[35] Centre hospitalier régional universitaire Bretonneau, Tours, France
[36] Université de Bordeaux, CRCM pédiatrique, center de Recherche Cardio-thoracique de Bordeaux, INSERM U1045, Bordeaux Imaging Center, CIC 1401 F-33000 Bordeaux, France
[37] Laboratory of Biochemistry, Hôpital Universitaire Necker Enfants Malades AP-HP Centre, Paris, France
[38] Pharmacie Delpech, Paris, France
[39] Université Paris Cité and Institut Cochin, Inserm U1016, Paris, France
[40] Respiratory Medicine and Cystic Fibrosis National Reference Center, Hôpital Cochin, AP-HP.Centre Université Paris Cité, Paris, France
[41] ERN-Lung CF network, Frankfurt, Germany
[42] Vaincre La Mucoviscidose, Paris, France
[43] Telethon Institute of Genetics and Medicine (TIGEM), Pozzuoli, Italy

[44] Centre de ressources et de compétences pour la mucoviscidose Adultes, Centre hospitalier régional universitaire de Nancy, Vandœuvre-Lès-Nancy, France
[45] Unité de Recherche Clinique, Hôpital Necker Enfants Malades, AP-HP, Paris, France
[46] Service de Médecine Génomique des Maladies de Système et d'Organe, Hôpital Cochin, Paris, France

### Methods

#### Plasmids and cell models

The cDNA of *CFTR* was subcloned into the pTracer expression vector as described by Fanen et al (1). CFTR mutants were generated by site-directed mutagenesis using the Quickchange II XL kit (Agilent, les Ulis, France). Fully sequenced clones were amplified and purified using Macherey Nagel Midipreps. Cells were transfected with lipofectamine 3000 (Invitrogen SARL). HEK293 cells were purchased from ATCC and cultivated in DMEM medium supplemented with 10% fetal calf serum.

#### Primary nasal epithelial cell sampling and culture

HNECs were cultured and expanded in serum-free PneumaCult Ex-Plus medium (StemCell Technologies, Vancouver, BC, Canada), supplemented with ROCK and SMAD inhibitors (Y-27632 and DMH-1, A-83-01 compounds). For the first days, culture medium also contained a mixture of different antibiotics (colistin, piperacillin, and tazobactam) to eradicate bacterial contamination. Differentiated epithelia were obtained by seeding human nasal epithelial cells (HNECs) at high density (500,000 cells/cm$^2$) on porous membranes (Snapwell inserts, code 3801, Corning Life Sciences, Corning, NY, USA). After 24 h, the medium was removed from both sides and replaced with Pneumacult ALI medium (StemCell Technologies, Vancouver, BC, Canada) on the basolateral side only. Epithelia differentiation (up to 16–18 days) was performed in air-liquid interface (ALI) condition (2–4).

#### Western Blot analysis

Samples were mixed with a Laemmli sample buffer and incubated at room temperature for 15 min, resolved using an 7% w/v acrylamide SDS-PAGE gel and transferred to nitrocellulose membranes. The membranes were then incubated in a blocking buffer (3% w/v BSA (bovine serum albumin) and 3% w/v milk in PBS-0.1% v/v Tween-20) for 1 h. Proteins of interest were immuno-detected with appropriate antibodies diluted in a 3% w/v BSA blocking buffer for 1 h.

CFTR was immuno-detected with mouse monoclonal (ab596, J.R. Riordan, University of North Carolina at Chapel Hill, and Cystic Fibrosis Foundation Therapeutics) diluted 1/1000 in blocking buffer. Finally, nitrocellulose membranes were incubated with appropriate secondary antibodies coupled to fluorochromes, as recommended by the manufacturer (Li-Cor, Bad Homburg, Germany). The detection of the WB results was performed with the Odyssey scanner (Li-Cor, Germany). Quantification of the WB was performed using ImageJ software. Lysates of primary nasal epithelia were generated following the previously described protocol (2, 3). In brief, to remove the mucus excess, the apical side of differentiated HNEC epithelia (ALI conditions for 16 days) were washed with warm HBSS containing 0.4% sodium bicarbonate for 3 h at 37 °C. After washing twice with warm complete PBS, the apical side of the filters was dried. Basolateral culture medium (Pneumacult ALI; StemCell Technologies, Vancouver, BC, Canada) was changed to treat cells with indicated correctors or vehicle (DMSO) for 24 h at 37 °C. The following day, the newly produced mucus was removed by washing the apical side of epithelia with warm HBSS 0.4% sodium bicarbonate at 37 °C for 30 min and then with warm complete PBS. Epithelia were lysed on ice by applying 100 µL/filter of ice-cold RIPA buffer (50 mM Tris-HCl pH 7.4, 150 mM NaCl, 1% Triton X-100, 0.5% Sodium deoxycholate, 0.1% SDS) plus proteases inhibitors (Merck KGaA, Darmstadt, Germany). Cell layers were scraped, collected in a tube, and left on ice for 15 min. To reduce the lysate viscosity, 5 × 22 G needle syringe passages followed by 5 × 27 G needle syringe passages were applied. Lysates were then cleared by centrifugation (15,000× g for 20 min at 4 °C). After centrifugation, the supernatant was transferred to a new tube and stored at −80 °C for subsequent analysis. The supernatant protein concentration was measured using a BCA assay (ThermoFisher Scientific,Waltham, MA, USA) following the manufacturer's instructions. Proteins (50 µg for HNEC epithelia) were separated on 4–15% gradient Criterion TGX Precast gels (Bio-rad Laboratories Inc., Hercules, CA, USA), transferred to a nitrocellulose membrane with a Trans-Blot Turbo system (Bio-rad Laboratories Inc., Hercules, CA, USA) and analyzed by Western blotting. CFTR was detected using the mouse monoclonal anti-CFTR (ab769, J.R. Riordan, University of North Carolina at Chapel Hill, and Cystic Fibrosis Foundation Therapeutics) while GAPDH was detected using the mouse monoclonal anti-GAPDH (sc-32233; Santa Cruz Biotechnology, Inc.), followed by either horseradish peroxidase (HRP)-conjugated anti-mouse IgG (ab97023; Abcam) or horseradish peroxidase (HRP)-conjugated goat anti-rabbit IgG (0031460; ThermoFisher Scientific, Waltham, MA, USA) and subsequently visualized by chemiluminescence using the SuperSignalWest Femto or West Dura Substrate (ThermoFisher Scientific,Waltham, MA, USA). Molecular Imager ChemiDoc XRS System (Bio-rad

Laboratories Inc., Hercules, CA, USA) was used to monitor the chemiluminescence. Images were analyzed with ImageJ software (National Institutes of Health, Bethesda). Profile analysis of different lanes was performed using the ImageJ software (National Institutes of Health, Bethesda, MD, USA). Impact of autophagy on CFTR biogenesis was investigated with two autophagy inhibitors, VPS inh (10 µM, 24 hours) and SAR-405 (2µM, 3 hours).

**Nano-Glo HiBiT complementation assays**

Total and surface CFTR quantification was performed using Nanoluciferase complementation assay (Nano-Glo® HiBiT Extracellular Detection System, Promega, France) enabled by the association of HiBiT tag inserted into the fourth extracellular loop of CFTR with its complementation partner Large BiT (LgBiT) [5]. The sequence encoding the eleven amino acid HiBiT peptide tag was inserted within the 4th extracellular loop of WT, L206W, F508del, or N1303K CFTR. HEK293 cells were transfected in six-well plates with Lipofectamine 3000 according the manufacturer's instructions (#L3000-015, Thermo Fisher Scientific, France) using 1,5 µg of pTracer plasmid, either empty or containing the cDNA encoding for WT or mutant CFTR. Transfected HEK-293 cells were plated onto white 96-well plates. The next day, cells were treated with ELX (3 µM) and TEZ (3 µM) for 24 hours. Cells were incubated with LgBit recombinant protein and nanoluciferase substrate, as per the manufacturer's instructions (#N2420, Nano-Glo® HiBiT Extracellular Detection System, Promega, France). After a 10-minute incubation at room temperature, luminescent signals corresponding to cell surface CFTR were acquired using a Mitras plate reader (Berthold, Thoiry, France). Cells were then permeabilized by adding NP-40 at a final concentration of 0.5%, enabling the diffusion of the LgBIT probe into the intracellular space for the quantification of the total CFTR. Background signals were subtracted (using empty pTracer), and quantification was performed for total CFTR level, membranar level and the cell surface ratio, defined as the cell surface CFTR signal normalized to the total CFTR signal.

To evaluate the stability of total CFTR protein, using the same experimental conditions, cells underwent when indicated a 24-hour treatment with ELX (3 µM) and TEZ (3 µM). Subsequently, protein synthesis was arrested by the addition of Cycloheximide (50 µg/mL, Sigma Aldrich Chimie SARL, France) at various time points. The cells were lysed directly in the presence of LgBit recombinant protein and nanoluciferase substrate, following the manufacturer's instructions (#N2420, Nano-Glo® HiBiT Lytic Detection System, Promega, France). The condition without cycloheximide, referred to as T0, serves as the baseline for normalizing each CFTR mutant or applied condition, thus corresponding to 100% of the protein.

**HS-YFP-based assay**

CFBE41o- cells having stable expression of the halide-sensitive yellow fluorescent protein (HS-YFP) were grown in MEM medium (Euroclone, Milan, Italy) supplemented with 10% FBS, 2 mM L-glutamine, 100 U/mL penicillin, and 100 µg/mL streptomycin (Euroclone, Milan, Italy). Vectors encoding WT-, N1303K- and F508del-CFTR variants were purchased from VectorBuilder (vector IDs available upon request; Neu-Isenburg, Germany). For the YFP assay, CFBE41o- cells (50,000 cells/well) stably expressing the halide-sensitive yellow fluorescent protein (HS-YFP) were reverse-transfected on clear-bottom 96-well black microplates (Corning Life Sciences, Corning, NY, USA) with 0.2 µg per well of the indicated vectors as previously described [2–4]. In brief, cells were transfected in Opti-MEM Reduced Serum Medium (ThermoFisher Scientific, Waltham, MA, USA) using Lipofectamine 2000 (ThermoFisher Scientific, Waltham, MA, USA) as the transfection agent. Opti-MEM was carefully replaced after 6 h, with culture medium without antibiotics. Twenty-four hours after transfection and plating, cells were treated with correctors or vehicle alone (DMSO) at the desired concentrations and incubated at 37 °C for an additional 24 h, prior to proceeding with the functional HS-YFP-based assay.

CFTR activity was determined by the HS-YFP microfluorimetric assay on CFBE41o-cells (transiently transfected to express different CFTR variants). Briefly, prior to the assay, CFBE41o- cells were washed with Dulbecco's PBS and then incubated for 25 min with 60 µL per well of Dulbecco's PBS containing forskolin (20 µM) and VX-770 (1 µM), at 37°C, to maximally stimulate the CFTR channel. Cells were then transferred to a microplate reader (Fluostar Optima; BMG Labtech, Offenburg, Germany), equipped with high-quality excitation (HQ500/20X: 500 ± 10 nm) and emission (HQ535/30M: 535±15 nm) filters for YFP (Chroma Technology, Bellows Falls, VT, USA). During the assay a continuous 14-s YFP fluorescence recording was performed, with 2 s before and 12 s after injection of 165 µL of an iodide-containing solution (Dulbecco's PBS where the NaCl was replaced by NaI; final $I^-$ concentration 100 mM). After subtracting the background, fluorescence data were normalized to the initial value. The $I^-$ influx rate was determined by fitting, for each well, the final 11 s of the data with an exponential function to extrapolate the initial slope (dF/dt). YFP quenching rate by I- enabled to assess CFTR activity.

**Interactomic studies**

Proximity labeling using TurboID [6, 7] identified proteins in the vicinity (<10nm) of WT-CFTR, F508del-CFTR and N1303K-CFTR. Independent experiments of HEK293 cells were transiently transfected for 48 h in 10 cm diameter poly-L-lysine coated dishes (n=3). Biotinylation was induced by adding biotin (Thermo Fisher Scientific, Illkirch-Graffenstaden, France) to the medium at 37 ◦C for 10 min (TurboID, 500 μM biotin).

NanoLC-MS/MS Protein Identification and Quantification S-TrapTM micro spin column (Protifi, Farmingdale, NY, USA) digestion was performed on streptavidin eluates in 4× Laemmli buffer according to the manufacturer's protocol but with 2 extra washing steps for thorough SDS elimination. Samples were digested with 2 µg of trypsin (Promega, Charbonnières-les-Bains, France) at 47 ◦C for 1 h 30 min. After elution, peptides were finally vacuum dried down and resuspended in 35 µL of 10% ACN and 0.1% TFA in HPLC-grade water prior to MS analysis. For each run, 5 µL was injected in a nanoRSLC-Q Exactive PLUS (RSLC Ultimate 3000) (Thermo Scientific, IllkirchGraffenstaden, France). Peptides were loaded onto a µ-precolumn (Acclaim PepMap 100 C18, cartridge, 300 µm i.d. × 5 mm, 5 µm) (Thermo Scientific, Illkirch-Graffenstaden, France) and were separated on a 50 cm reversed-phase liquid chromatographic column (0.075 mm ID, Acclaim PepMap 100, C18, 2 µm) (Thermo Scientific, Illkirch-Graffenstaden, France). The chromatography solvents were (A) 0.1% formic acid in water and (B) 80% acetonitrile, 0.08% formic acid. Peptides were eluted from the column with the following gradients: 5% to 40% B (120 min), 40% to 80% (1 min). At 121 min, the gradient stayed at 80% for 5 min, and at 127 min, it returned to 5% to re-equilibrate the column for 20 min before the next injection. One blank was run between each series to prevent sample carryover. Peptides eluting from the column were analyzed by data-dependent MS/MS, using the top-10 acquisition method. Peptides were fragmented using higher-energy collisional dissociation (HCD). Briefly, the instrument settings were as follows: the resolution was set to 70,000 for MS scans and 17,500 for the data-dependent MS/MS scans in order to increase speed. The MS AGC target was set to 3.106 counts with a maximum injection time set to 200 ms, while the MS/MS AGC target was set to 1.105 with a maximum injection time set to 120 ms. The MS scan range was from 400 to 2000 m/z. 4.7.

The MS files were processed with the MaxQuant software version 2.0.1.0 and searched with the Andromeda search engine against the database of Homo sapiens from Swiss-Prot 04/2020. To search for parent mass and fragment ions, we set an initial mass deviation of 4.5 ppm and 20 ppm, respectively. The minimum peptide length was set to 7 amino acids, and strict specificity for trypsin cleavage was required, allowing up to two missed cleavage sites.

Carbamidomethylation (Cys) was set as fixed modification, whereas oxidation (Met) and N-term acetylation were set as variable modifications. A match between the runs was allowed. LFQ minimum ratio count was set to 2. The false discovery rates (FDR) at the protein and peptide levels were set to 1%. Scores were calculated in MaxQuant, as described previously (7). The reverse and common contaminants hits were removed from the MaxQuant output. Proteins were quantified according to the MaxQuant label-free algorithm using LFQ intensities (8). Fragmentation visualization spectra were also extracted using the MQviewer integrated in the Maxquant software. LFQ intensities were used as input in MassDynamics tools to perform pairwise differential expression analysis between CFTR mutants and the WT condition. Volcano plot shows statistical significance (P value) versus magnitude of change (log2 fold change). Over Representation Analysis (ORA) tests whether any pathway in the Reactome database (9, 10) is significantly over-represented in proteins enriched in N1303K-CFTR compared to F508del-CFTR ($Log_2$ Fold Change >1). False discovery rates are calculated using the Benjamini–Hochberg procedure. Enrichment Bar plot showing the top 10 over-represented pathways ordered by -log10 estimated False Discovery Rate (FDR).

MaxQuant protein LFQ intensities were used to asses a probabilistically scoring protein-protein interaction of CFTR using SAINTexpress (11) (https://reprint-apms.org/). Knowed CFTR partners from CyFi-map (12) were used to evaluate N1303K proximal interactome. Average intensities protein identifies with TurboID for CFTR-mutants and BFDR evaluating protein-protein interaction are visualize with Dot plot that have been generated using ProHitz-viz tools (13) (https://prohits-viz.lunenfeld.ca/).

**Modeling of protein inhibition synthesis by cycloheximide**

The inhibition of protein synthesis by cycloheximide over time was analysed using the nonlinear modelling software program Monolix version 2023R1 (http://lixoft.com). A typical plateau function, as $f(t) = 100 \times (1 - \frac{t}{t_{50} + t})$ was used to estimate the time to reach 50% of inhibition ($t_{50}$). Covariates cross-effect of N1303K-CFTR, F508del-CFTR and ELX/TEZ were tested on $t_{50}$ parameter. Overall parameters were estimated by the Markov Chain Monte Carlo-Stochastic Approximation Expectation Maximization algorithm (MCMC-SAEM).

Overall structural model parameters were accurately estimated given their relative standard errors of estimates (RSE% < 30%). The covariates cross-effect of N1303K-CFTR, F508del-CFTR and ELX/TEZ were significant on $t_{50}$ parameter ($p<0.05$, Wald test).

**Interactomic studies**

Proximity labeling assays identified proteins in the vicinity (<10nm) of WT or mutated CFTR. Compared to F508dek, N1303K-CFTR were found in close proximity with proteins from the Golgi apparatus, endosomes and plasma membrane (**Figure 2f and 2g**). Analysis of known WT-CFTR partners as reported in CyFi-map (12) showed that N1303K-CFTR interactome included significantly more partners from the sorting/recycling endosome of CFTR (e.g; COPII sorting complex – SEC23A and SAR1B ; SNARE protein - VAMP8; Syntaxins - STX6 and STX7; Synaptosomal-associated protein - SNAP23; retromer complex proteins - SNX27) and membranar or submembranar proteins (e.g; Filamin A – FLNA; ERM complex - EZR; Lemur Tyrosine Kinase 2 - LMTK2; NHERF family proteins - SLC9A3R1 and SLC9A3R2; clatherin triskelion - AP2B1 and AP2B2) (**Figure Supplemental 5a**).

N1303K-CFTR is also in close proximity with Endoplasmic Reticulum proteins according CyFi map. This is similar to ER interactome of F508del-CFTR including ER folding and quality control (**Figure Supplemental 5b**) (14) in contrast to WT-CFTR. Regarding proximal interaction with proteasomal complex proteins, N1303K and F508del share a similar profile albeit the relative abundance is decrease in N1303K compared to F508del-CFTR (**Figure Supplemental 5c**).

**Italian pwCF**

Twenty-eight patients compound heterozygous for N1303K and a MF, non-rescuable *CFTR* variant resembling a null allele (donor IDs: GE010, GE011, GE018, FI034, FI047, FI053, FI057, FI064, GE068, ME088, ME090, ME092, ME094, ME119, GE132, FI134, GE144, GE156, ME163, GE226, MI240, MI247, MI248, MI258, MI259, MI264, GE326 and GE331) and five patients homozygous for N1303K (ME084, ME087, GE207, MI239, MI291) were included in this study. One healthy subject (donor ID: Ctr069), four patients homozygous for F508del (donor IDs:TT003, TT005, AN232, AN235, and AN237), one patient heterozygous for F508del (donor ID: AN238), three patients compound heterozygous for L206W and a MF, non-rescuable *CFTR* variant (GE030, FI051, and ME169), three patients compound heterozygous for R334W and a MF, non-rescuable *CFTR* variant (GE016, FI058, and GE211) and one patient homozygous for the G542X (donor ID: GE220) were enrolled as further controls.

**Genetic studies**

Extensive genetic screening was performed by next generation sequencing in 5 patients showing a mild or transient response to treatment (patient # 6, 8, 9, 15,16), to search for: i) *CFTR* complex alleles in *cis* with N1303K, including variants resulting in different haplotypes (c. 744-33GATT6/7, c.869+11C/T) [15]; ii) variants in other genes that were shown to modify airway surface liquid homeostasis *TMEM16A, SLC26A4, ATP12A, and SCNN1A/B/G* and *SERPINA1* gene, involved in alpha1-antitrypsin deficiency [16].
Only the frequent haplotype described in cis with N1303K (c.[744-33GATT[6];869+11C>T]) was evidenced (15) and no other *CFTR* variant was identified as a complex allele. No correlation could thus be made with the treatment response. Indeed, both the haplotype in *cis* with N1303K and the reference WT haplotype induce minor exon 7 skipping (7% in minigenes) (15, 17) which suggests that they should not affect either the baseline CFTR function or the response to ETI.
We found in patient #16, a heterozygous *SCNN1A* variant, p.Arg181Trp, which increases ENaC activity by 1.75 fold (18). Such an effect was suggested in the patient by the response to amiloride at -56.5 µA/cm$^2$, 1.5-fold of that observed in other HNECs non-F508del MF variants. Patient #15 was heterozygous for a *SLC26A4* pathogenic variant involved in Pendred syndrome, an autosomal recessive disease associating hearing loss and thyroid iodine organification defect (19). Patient #6 was compound heterozygous for Z and S *SERPINA1* variants, a genotype responsible for alpha1-antitrypsin deficiency.

For two of the N1303K homozygous patients (ME084 and ME087, displaying different drug respectively responsiveness (respectively weak and best response), transcript analysis was performed to exclude additional coding variants and unconventional splicing events. We did not identify any additional coding variants in cis with N1303K that could affect gene expression levels, pre-mRNA splicing or transcript stability. Moreover, the genotype of the M470V polymorphism was also defined in these patients to exclude any correlation with drug response.

**Bibliography**


1. Fanen P, Clain J, Labarthe R, Hulin P, Girodon E, Pagesy P, Goossens M, Edelman A. Structure-function analysis of a double-mutant cystic fibrosis transmembrane conductance regulator protein occurring in disorders related to cystic fibrosis. *FEBS Lett* 1999; 452: 371–374.

2. Tomati V, Costa S, Capurro V, Pesce E, Pastorino C, Lena M, Sondo E, Di Duca M, Cresta F, Cristadoro S, Zara F, Galietta LJV, Bocciardi R, Castellani C, Lucanto MC, Pedemonte N. Rescue by elexacaftor-tezacaftor-ivacaftor of the G1244E cystic fibrosis mutation’s stability and gating defects are dependent on cell background. *J Cyst Fibros* 2023; 22: 525–537.

3. Terlizzi V, Pesce E, Capurro V, Tomati V, Lena M, Pastorino C, Bocciardi R, Zara F, Centrone C, Taccetti G, Castellani C, Pedemonte N. Clinical Consequences and Functional Impact of the Rare S737F CFTR Variant and Its Responsiveness to CFTR Modulators. *Int J Mol Sci* 2023; 24: 6576.

4. Sondo E, Cresta F, Pastorino C, Tomati V, Capurro V, Pesce E, Lena M, Iacomino M, Baffico AM, Coviello D, Bandiera T, Zara F, Galietta LJV, Bocciardi R, Castellani C, Pedemonte N. The L467F-F508del Complex Allele Hampers Pharmacological Rescue of Mutant CFTR by Elexacaftor/Tezacaftor/Ivacaftor in Cystic Fibrosis Patients: The Value of the Ex Vivo Nasal Epithelial Model to Address Non-Responders to CFTR-Modulating Drugs. *Int J Mol Sci* 2022; 23: 3175.

5. Baatallah N, Elbahnsi A, Chevalier B, Castanier S, Mornon J-P, Pranke I, Edelman A, Sermet-Gaudelus I, Callebaut I, Hinzpeter A. Acting on the CFTR Membrane-Spanning Domains Interface Rescues Some Misfolded Mutants. *Int J Mol Sci* 2022; 23: 16225.

6. Branon TC, Bosch JA, Sanchez AD, Udeshi ND, Svinkina T, Carr SA, Feldman JL, Perrimon N, Ting AY. Efficient proximity labeling in living cells and organisms with TurboID. *Nat Biotechnol* 2018; 36: 880–887.

7. Chevalier B, Baatallah N, Najm M, Castanier S, Jung V, Pranke I, Golec A, Stoven V, Marullo S, Antigny F, Guerrera IC, Sermet-Gaudelus I, Edelman A, Hinzpeter A. Differential CFTR-Interactome Proximity Labeling Procedures Identify Enrichment in Multiple SLC Transporters. *IJMS* 2022; 23: 8937.

8. Cox J, Mann M. MaxQuant enables high peptide identification rates, individualized p.p.b.-range mass accuracies and proteome-wide protein quantification. *Nat Biotechnol* 2008; 26: 1367–1372.

9. Wu G, Haw R. Functional Interaction Network Construction and Analysis for Disease Discovery. *Methods Mol Biol* 2017; 1558: 235–253.

10. Joshi-Tope G, Gillespie M, Vastrik I, D’Eustachio P, Schmidt E, de Bono B, Jassal B, Gopinath GR, Wu GR, Matthews L, Lewis S, Birney E, Stein L. Reactome: a knowledgebase of biological pathways. *Nucleic Acids Res* 2005; 33: D428-432.

11. Teo G, Liu G, Zhang J, Nesvizhskii AI, Gingras A-C, Choi H. SAINTexpress: Improvements and additional features in Significance Analysis of INTeractome software. *Journal of Proteomics* 2014; 100: 37–43.

12. Pereira C, Mazein A, Farinha CM, Gray MA, Kunzelmann K, Ostaszewski M, Balaur I, Amaral MD, Falcao AO. CyFi-MAP: an interactive pathway-based resource for cystic fibrosis. *Sci Rep* 2021; 11: 22223.

13. Knight JDR, Choi H, Gupta GD, Pelletier L, Raught B, Nesvizhskii AI, Gingras A-C. ProHits-viz: a suite of web tools for visualizing interaction proteomics data. *Nat Methods* 2017; 14: 645–646.

14. Farinha CM, Canato S. From the endoplasmic reticulum to the plasma membrane: mechanisms of CFTR folding and trafficking. *Cell Mol Life Sci* 2017; 74: 39–55.

15. Farhat R, Puissesseau G, El-Seedy A, Pasquet M-C, Adolphe C, Corbani S, Megarbané A, Kitzis A, Ladeveze V. N1303K (c.3909C>G) mutation and splicing: implication of its c.[744-33GATT(6); 869+11C>T] complex allele in CFTR exon 7 aberrant splicing. *Biomed Res Int* 2015; 2015: 138103.

16. Zajac M, Dreano E, Edwards A, Planelles G, Sermet-Gaudelus I. Airway Surface Liquid pH Regulation in Airway Epithelium Current Understandings and Gaps in Knowledge. *Int J Mol Sci* 2021; 22: 3384.

17. Farhat R, El-Seedy A, Pasquet M-C, Corbani S, Megarbané A, Kitzis A, Ladeveze V. Three Complex alleles associated with N1303K mutation and their molecular consequences. *Cell Mol Biol (Noisy-le-grand)* 2022; 68: 52–59.

18. Azad AK, Rauh R, Vermeulen F, Jaspers M, Korbmacher J, Boissier B, Bassinet L, Fichou Y, des Georges M, Stanke F, De Boeck K, Dupont L, Balascáková M, Hjelte L, Lebecque P, Radojkovic D, Castellani C, Schwartz M, Stuhrmann M, Schwarz M, Skalicka V, de Monestrol I, Girodon E, Férec C, Claustres M, Tümmler B, Cassiman J-J, Korbmacher C, Cuppens H. Mutations in the amiloride-sensitive epithelial sodium channel in patients with cystic fibrosis-like disease. *Hum Mutat* 2009; 30: 1093–1103.

19. Ito T, Muskett J, Chattaraj P, Choi BY, Lee KY, Zalewski CK, King KA, Li X, Wangemann P, Shawker T, Brewer CC, Alper SL, Griffith AJ. SLC26A4 mutation testing for hearing loss associated with enlargement of the vestibular aqueduct. *World J Otorhinolaryngol* 2013; 3: 26–34.
