## Supplemental table for "Beyond Kaftrio : mechanistic insights to maximize N1303K-CFTR rescue"

**Supplemental Table 1. Genotype of the Italian subjects**

| Donor ID | Allele 1 | Allele 2 |
|---|---|---|
| ME084 | N1303K | N1303K |
| ME087 | N1303K | N1303K |
| GE207 | N1303K | N1303K |
| MI239 | N1303K | N1303K |
| MI291 | N1303K | N1303K |
| GE010 | N1303K | 394delTT |
| GE011 | N1303K | 1717-1G>A |
| GE018 | N1303K | 711+1G>T |
| FI034 | N1303K | G542X |
| FI047 | N1303K | 1717-1G>A |
| FI053 | N1303K | Delexon22-24 |
| FI057 | N1303K | E585X |
| FI064 | N1303K | 1717-1G>A |
| GE068 | N1303K | W1282X |
| ME088 | N1303K | R553X |
| ME090 | N1303K | I507del |
| ME092 | N1303K | G542X |
| ME094 | N1303K | G542X |
| ME119 | N1303K | 3659delC |
| GE132 | N1303K | 2183AA>G |
| FI134 | N1303K | 1717-1G>A |
| GE144 | N1303K | G542X |
| GE156 | N1303K | 711+1G>T |
| ME163 | N1303K | 1002-1113_1002-1110delGAAT |
| GE226 | N1303K | 1713del |
| MI240 | N1303K | L732X |
| MI247 | N1303K | G542X |
| MI248 | N1303K | 4016insT |
| MI258 | N1303K | 711+5G>A |
| MI259 | N1303K | C276X |
| MI264 | N1303K | 852del22bp |
| GE331 | N1303K | 1717-1G>A |
| GE326 | N1303K | G542X |
| TT003 | F508del | F508del |
| TT005 | F508del | F508del |
| AN232 | F508del | F508del |
| AN235 | F508del | F508del |
| AN238 | F508del | 2183AA>G |
| GE030 | L206W | G542X |
| FI051 | L206W | 3659delC |
| ME169 | L206W | 2183AA>G |
| GE016 | R334W | G542X |
| FI058 | R334W | 1711_1716+11del |
| GE211 | R334W | 2183AA>G |
| GE220 | G542X | G542X |
| Ctr069 | - | - |

**Supplemental Table 2. Clinical data of pwCF carrying the N1303K variant treated with ETI**

| Patient | Data origin | CFTR variant | CFTR variant | Sweat chloride (mmol/l) | | Percent predicted $FEV_1$ | |
|---|---|---|---|---|---|---|---|
| | | Allele 1 | Allele 2 | Baseline | ETI | Baseline | ETI |
| **1** | France (14,18) | N1303K | N1303K | 109 | 87 | 19 | 30 |
| **2** | France (14,18) | N1303K | N1303K | 105 | 96 | 33 | 92 |
| **3** | France (14,18) | N1303K | N1303K | 93 | 92 | 44 | 69 |
| **4** | France (14,18) | N1303K | N1303K | 114 | 76 | 23 | 32 |
| **5** | France (14,18) | N1303K | N1303K | 96 | 91 | 20 | 30 |
| **6*** | **France (14,18)** | **N1303K** | **N1303K** | N/A | N/A | 23 | 34 |
| **7** | France (14,18) | N1303K | N1303K | N/A | 84 | 48 | 71 |
| **8** | France (21) | N1303K | N1303K | 103 | 99 | 79 | 82 |
| **9** | **France (unpublished)** | **N1303K** | **N1303K** | 100 | 93 | 83 | 87 |
| **10*** | **France (unpublished** | **N1303K** | **N1303K** | 88 | 86 | 56 | 69 |
| **11** | France (unpublished | N1303K | N1303K | 110 | 112 | 68 | 85 |
| **12** | France (unpublished | N1303K | N1303K | 95 | 96 | 69 | 95 |
| **13** | France (unpublished | N1303K | N1303K | 94 | 94 | 100 | 117 |
| **14** | France (unpublished | N1303K | N1303K | 103 | 90 | 129 | 132 |
| **15** | France (unpublished | N1303K | N1303K | 97 | 84 | 88 | 100 |
| **16** | Israel (17) | N1303K | N1303K | 111 | 80 | 90 | 109 |
| **17** | Israel (17) | N1303K | N1303K | 98 | 88 | 47 | 65 |
| **18** | Germany (19) | N1303K | N1303K | 99 | 68 | 54 | 64 |
| **19** | France (14,18) | N1303K | R1162X | 99 | 90 | 32 | 61 |
| **20*** | **France (14,18)** | **N1303K** | **R1162X** | 131 | 95 | 46 | 54 |
| **21** | **France (18)** | **N1303K** | **G542X** | 104 | 96 | 26 | 27 |
| **22** | France (18) | N1303K | del3-10,14b-16 | 102 | 94 | 48 | 73 |
| **23** | France (21) | N1303K | H199R | 95 | 75 | 40 | 60 |
| **24** | France (21) | N1303K | 2789+5G>A | 100 | 71 | N/A** | N/A** |
| **25** | France (21) | N1303K | 1898+5G>A | 106 | 68 | 50 | 53 |
| **26** | France (21) | N1303K | 4326delTC | 108 | 90 | 92 | 128 |
| **27** | France (21) | N1303K | M1V | 106 | 105 | 43 | 64 |
| **28** | France (21) | N1303K | 711+1G>T | 109 | 140 | 63 | 73 |
| **29** | France (21) | N1303K | E585X | 118 | 89 | 53 | 82 |
| **30** | France (21) | N1303K | R553X | 91 | 88 | 41 | 58 |
| **31** | France (21) | N1303K | R553X | 107 | 103 | 50 | 60 |
| **32** | France (21) | N1303K | W1282X | 109 | 103 | 49 | 65 |
| **33** | France (21) | N1303K | 3120+1G>A | 110 | 101 | 61 | 101 |
| **34** | France (21) | N1303K | 3659delC | 101 | 84 | 67 | 80 |
| **35** | Israel (17) | N1303K | W1282X | 130 | 118 | 81 | 111 |
| **36** | Israel (17) | N1303K | E819X | 115 | 110 | 97 | 114 |
| **37** | Israel (17) | N1303K | W1282X | 119 | 112 | 30 | 39 |
| **38** | Israel (17) | N1303K | W1282X | 107 | 114 | 68 | 86 |
| **39** | Israel (17) | N1303K | 3121-1G>A | 114 | 114 | 85 | 111 |
| **40** | Israel (17) | N1303K | G542X | 96 | 95 | 56 | 66 |
| **41** | USA (15) | N1303K | E193X | 108 | 95 | 87 | 108* |
| **42** | USA (22) | N1303K | Q493X | 115 | 113 | 84 | 93 |
| **43** | France (unpublished | N1303K | 3600G>A | 60 | 100 | 78 | 90 |
| **44** | France (unpublished | N1303K | CFTRdele4-21 | 104 | 95 | 61 | 111 |
| **45** | France (unpublished | N1303K | I507del | 95 | 87 | 83 | 97 |
| **46** | France (unpublished | N1303K | 1677delTA | 107 | 105 | 95 | 101 |
| **47** | France (unpublished | N1303K | Y1092X | 120 | 113 | 62 | 70 |
| **48** | France (unpublished | N1303K | 1717-1G>A | - | 85 | 76 | 79 |

| | | | | | | | |
|---|---|---|---|---|---|---|---|
| **49** | France (unpublished | N1303K | 1717-1G>A | 98 | 90 | 92 | 110 |
| **50** | France (unpublished | N1303K | Q290X | 143 | 120 | 90 | 98 |
| **51** | France (unpublished | N1303K | 711+1G>T | 104 | 85 | 32 | 38 |
| **52** | France (unpublished | N1303K | G542X | 99 | 99 | 62 | 90 |
| **53** | France (unpublished | N1303K | Q493X | 112 | 106 | 58 | 95 |
| **54** | France (unpublished | N1303K | 3120+1G>A | - | 101 | 61 | 101 |
| **55** | France (unpublished | N1303K | 2640delT | 100 | 91 | 62 | 75 |
| **56** | France (unpublished | N1303K | 2789+5G>A | 86 | 81 | 79 | 92 |
| **57** | France (unpublished | N1303K | 1811+1,6kbA>G | 100 | 98 | 64 | 78 |
| **58** | France (unpublished | N1303K | G542X | 110 | 90 | 112 | 132 |
| **59** | France (unpublished | N1303K | 2789+5G>A | 85 | 67 | 56 | 67 |
| **60** | France (unpublished | N1303K | 4374+1G>A | 92 | 97 | 68 | 84 |
| **61** | France (unpublished | N1303K | G542X | 94 | 117 | 113 | 130 |
| **62** | France (unpublished | N1303K | G542X | 110 | 90 | 112 | 132 |
| **63** | France (unpublished | N1303K | 2622+1G>A | 110 | 102 | 110 | 95 |
| **64** | France (unpublished | N1303K | G542X | 94 | 113 | 136 | 130 |
| **65** | France (unpublished | N1303K | 2622+1G>A | 110 | 102 | 110 | 95 |

*- patients who experienced a respiratory degradation after initial improvement

Patients 6, 9, 10, 20, 21 - sequencing
